# Activation of TIR signaling is required for pattern-triggered immunity

**DOI:** 10.1101/2020.12.27.424494

**Authors:** Hainan Tian, Siyu Chen, Zhongshou Wu, Kevin Ao, Hoda Yaghmaiean, Tongjun Sun, Weijie Huang, Fang Xu, Yanjun Zhang, Shucai Wang, Xin Li, Yuelin Zhang

## Abstract

Plant immune responses are mainly activated by two types of receptors. Plasma membrane-localized pattern recognition receptors (PRRs) recognize conserved features of microbes, and intracellular nucleotide-binding leucine rich repeat receptors (NLRs) recognize effector proteins from pathogens. NLRs possessing N-terminal Toll/interleukin-1 receptor (TIR) domains (TNLs) activate two parallel signaling pathways via the EDS1/PAD4/ADR1s and the EDS1/SAG101/NRG1s modules. The relationship between PRR-mediated pattern-triggered immunity (PTI) and TIR signaling is unclear. Here we report that activation of TIR signaling plays a key role in PTI. Blocking TIR signaling by knocking out components of the EDS1/PAD4/ADR1s and EDS1/SAG101/NRG1s modules results in attenuated PTI responses such as reduced salicylic acid (SA) levels and expression of defense genes, and compromised resistance against pathogens. Consistently, PTI is attenuated in transgenic plants that have reduced accumulation of NLRs. Upon treatment with PTI elicitors such as flg22 and nlp20, a large number of genes encoding TNLs or TIR domain-containing proteins are rapidly induced, likely responsible for activating TIR signaling during PTI. In support, overexpression of some of these genes results in activation of defense responses. Overall, our study reveals that TIR signaling activation is an important mechanism for boosting plant defense during PTI.

## Introduction

Immune receptors are essential for non-self recognition and defense activation in multicellular organisms^1,2^. Plants use pattern recognition receptors (PRRs), which include transmembrane receptor-like kinases (RLKs) and receptor-like proteins (RLPs), to detect conserved components of microbes collectively known as pathogen-associated molecular patterns (PAMPs), and activate pattern-triggered immunity (PTI)^3^. Unlike RLKs, RLPs do not have a cytoplasmic kinase domain and usually transduce defense signals through adaptor RLKs such as BRI1-ASSOCIATED RECEPTOR KINASE 1 (BAK1) and SUPPRESSOR OF BIR1 1 (SOBIR1)^4^. As an example, RLK FLAGELLIN-SENSITIVE 2 (FLS2) recognizes flg22, a conserved peptide from bacterial flagellin^5,6^. In contrast, *Arabidopsis* RLP23 recognizes a 20-amino-acid motif (nlp20) widely found in most NECROSIS AND ETHYLENE-INDUCING PEPTIDE 1-LIKE PROTEINS (NLPs) of microbes^7,8^. RLP23 constitutively associates with SOBIR1 and binding of nlp20 induces formation of a tripartite complex consisting RLP23, SOBIR1, and BAK1, leading to activation of downstream immune signaling^8^.

Activation of PTI typically leads to production of reactive oxygen species (ROS), activation of MITOGEN-ACTIVATED PROTEIN KINASES (MAPKs), increased biosynthesis of the defense hormone salicylic acid (SA) and up-regulation of defense-related genes^9^. Receptor-like cytoplasmic kinases (RLCKs), which have a kinase domain similar to RLKs but lack a transmembrane motif and extracellular ligand-binding domain, play crucial roles in transducing defense signals downstream of PRRs^10^.

To promote virulence, microbial pathogens deliver a variety of effector proteins to interfere with PTI and facilitate nutrient acquisition from plants^11^. Recognition of these pathogen effectors by plant immune receptors leads to the activation of effector-triggered immunity (ETI). The majority of intracellular nucleotide-binding leucine-rich repeat proteins (NLRs) serve as sensors for effectors^12^. These sensor NLRs (sNLRs) with an N-terminal coiled-coil (CC) domain or a Toll/interleukin-1 receptor (TIR) domain are knowns as CNLs and TNLs, respectively. Distinct mechanisms are used by the CC and TIR domains to activate defense signaling. The CC domain of HOPZ-ACTIVATED RESISTANCE 1 (ZAR1) was suggested to form a narrow pore on the plasma membrane to trigger cell death and plant immunity^13^. On the other hand, the TIR domains of many TNLs were shown to possess nicotinamide adenine dinucleotide (NAD+) hydrolase (NADase) activity, which is required for activation of downstream immune responses^14,15^. Intriguingly, two small groups of helper NLRs (hNLRs) in the ADR1 and NRG1 family, which carry an N-terminal RESISTANCE TO POWDERY MILDEW 8 (RPW8)-like CC (CC_R_) domain, function downstream of TNLs^16–21^. ADR1s play a critical role in activating SA biosynthesis^22^, while NRG1s are required for TNL-induced cell death^17,19^. In addition, three related lipase-like proteins, EDS1 (ENHANCED DISEASE SUSCEPTIBILITY 1)/PAD4 (PHYTOALEXIN DEFICIENT 4)/SAG101 (SENSECENCE-ASSOCIATED GENE 101), also function downstream of TNLs^23^. EDS1 form distinct protein complexes with PAD4 or SAG101. The EDS1/PAD4 complex functions in the same defense pathway as ADR1s, whereas the EDS1/SAG101 complex works together with NRG1 to promote cell death^17,20^.

SA plays diverse and critical roles in plant immunity^24,25^. It is required for PTI and ETI in local infection sites, as well as systemic acquired resistance (SAR), which confers protection against secondary infections in distal tissues. In *Arabidopsis*, pathogen-induced SA is mainly synthesized from isochorismate, which is produced from chorismate by ISOCHORISMATE SYNTHASE 1 (ICS1)^26^. PBS3 catalyzes the next step of conjugation of glutamate to isochorismate, and the resulting isochorismate-9-Glu subsequently decomposes to produce SA^27,28^. SAR-DEFICIENT 1 (SARD1) and CALMODULIN BINDING PROTEIN 60g (CBP60g) are two major transcription factors regulating SA biosynthetic genes during pathogen infection^29,30^. Increased *SARD1* expression and SA accumulation are two early events downstream of PRR activation during PTI^9^.

NLR homeostasis control is essential for regulating ETI immune output. Ubiquitination plays crucial roles in regulating the NLR protein levels. For example, the turnover of the TNL SNC1 (SUPPRESOR OF *npr1*, CONSTITUTIVE 1) and CNLs RPS2 (RESISTANCE TO *Pseudomonas syringae* 2) and SUMM2 (SUPPRESOR OF *mkk1 mkk2*, 2) is controlled by the Skp, Cullin, F-box (SCF) E3 ligase SCF^CPR131,32^. Two other E3 ligases MUSE1 (MUTANT, *snc1*-ENHANCING 1) and MUSE2 promote the degradation of several TNLs which pair with SNC1^33^. Another E3 ligase UBR7 interacts with tobacco TNL N to control its levels^34^. Recently it was also shown that the homeostasis of sensor NLRs is broadly regulated by the redundant E3 ligases SNIPER1 (*snc1*-INFLUENCING PLANT E3 LIGASE REVERSE GENETIC SCREEN) and SNIPER2^35^. Overexpression of *SNIPER1* leads to globally reduced sNLR levels and attenuated ETI responses.

Immune signaling mediated by PRRs and NLRs has been studied separately in the past; the connection between them is rarely explored. In this study, we tested PAMP-induced responses in *Arabidopsis SNIPER1* overexpression lines and mutants deficient in TNL signaling. Inhibition of NLR accumulation or abolishment of TIR signaling resulted in reduced SA accumulation and compromised PTI, suggesting that activation of TIR signaling plays a crucial role in PTI.

## Results

### PTI is compromised in *SNIPER1* overexpression lines

Overexpression of *SNIPER1* leads to reduced accumulation of sNLRs and compromised ETI^35^. To test whether PTI is affected in *SNIPER1* overexpression lines, we challenged them with *Pseudomonas syringae pv. tomato* (*Pto*) DC3000 *hrcC*, a bacterial strain unable to secret effectors due to a defect in the type III secretion system. Growth of *Pto* DC3000 *hrcC* was significantly higher in the *SNIPER1* overexpression lines than in wild type (WT) plants (Figure 1A). As no effectors can be delivered into host cells by *Pto* DC3000 *hrcC*, enhanced growth of this strain in the *SNIPER1* overexpression lines suggests a PTI deficiency. Next, we examined if other PTI responses are affected in these *SNIPER1* overexpression lines. Two defense-related genes *SARD1* and *FMO1* (*FLAVIN-CONTAINING MONOOXYGENASES*) are quickly induced upon infection. The expression levels of *SARD1* and *FMO1* after infection by *Pto* DC3000 *hrcC* were significantly reduced in the *SNIPER1* overexpression lines compared to WT (Figure 1B-C). Furthermore, we measured the SA accumulation induced by *Pto* DC3000 *hrcC*. Both free and glucose-conjugated SA (SAG) levels in the *SNIPER1* overexpression lines were significantly lower than those in the WT plants upon *Pto* DC3000 *hrcC* treatment (Figure 1D, S1A). Taken together, these data suggest that a general reduction of sNLR accumulation due to *SNIPER1* overexpression leads to compromised PTI.

**Figure 1.**
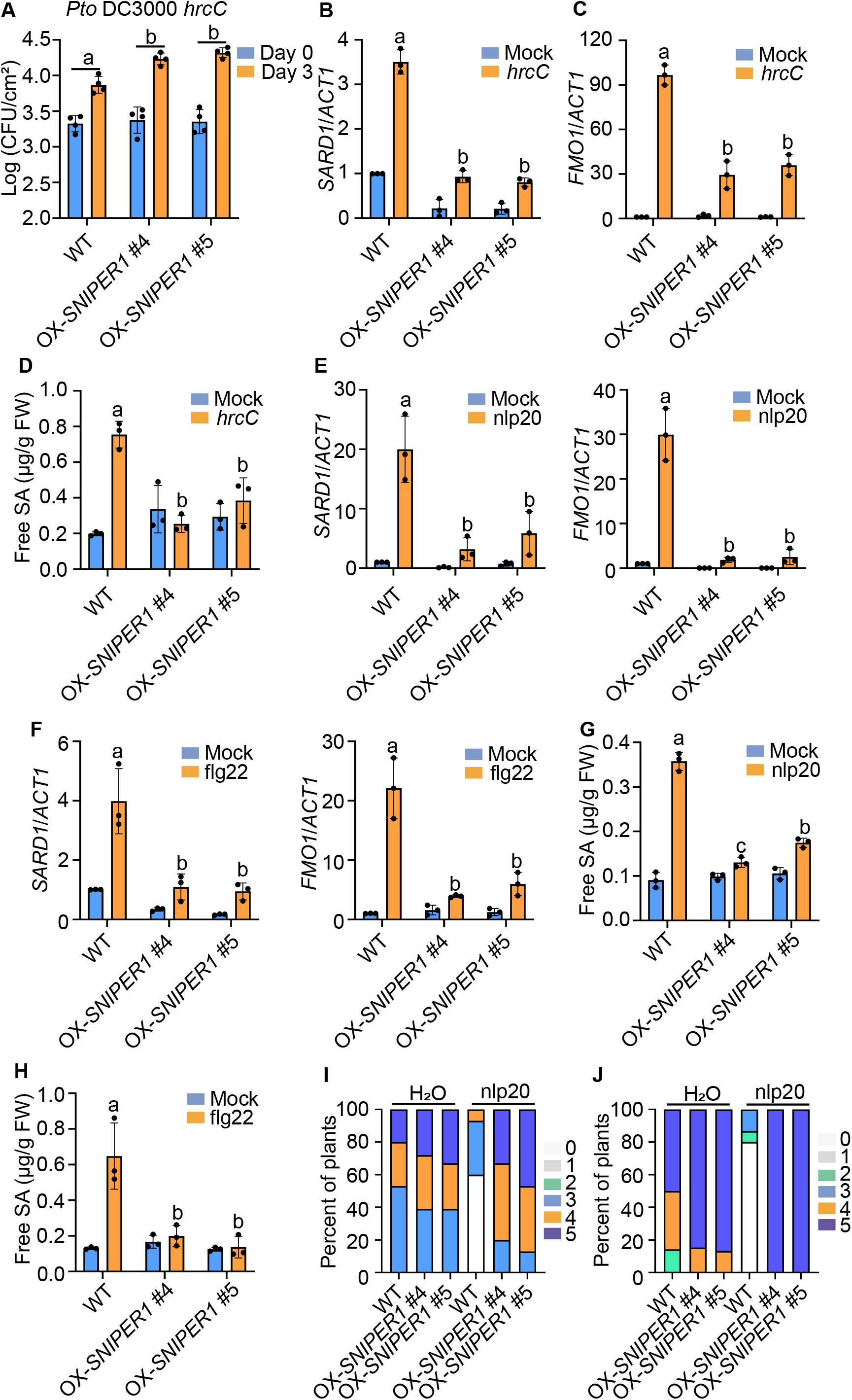
Overexpression of *SNIPER1* leads to attenuation of *Pto* DC3000 *hrcC*, flg22 and nlp20-induced immunity. (A) Growth of *Pto* DC3000 *hrcC* in wild type (WT) Col-0 and two independent *SNIPER1* overexpression *(OX-SNIPER1)* lines. Leaf discs were collected 0 days (Day 0) or 3 days (Day 3) after bacterial infiltration (OD_600_=0.002). Error bars represent standard deviation (SD) from four biological replicates. The growth of *Pto* DC3000 *hrcC* in different genotypes on Day 3 was compared using two-way ANOVA test, and different letters indicate genotypes with statistical differences (*p* < 0.05; n = 4). The experiment was repeated twice with similar results. (B, C) Relative expression levels of *SARD1* (B) and *FMO1* (C) in WT and *OX-SNIPER1* plants upon *Pto* DC3000 *hrcC* infection. Total RNA was isolated from leaf tissues of 25-d-old soil-grown plants 12 hours after infiltration with *Pto* DC3000 *hrcC* (OD_600_=0.05) or 10 mM MgCl2 (Mock). qPCR was used to examine the genes expression levels. *ACT1* was used for normalization, and the expression of each gene in mock-treated WT plants was set as 1. Error bars represent SD from three different biological replicates. Different letters indicate genotypes with statistical differences (*p* < 0.05, one-way ANOVA test; n=3). The experiment was repeated twice with similar results. (D) Free salicylic acid (SA) levels in four-week-old WT and OX-*SNIPER1* plants 12 hours after treatment with 10 mM MgCl2 (mock) or *Pto* DC3000 *hrcC*. Different letters indicate genotypes with statistical differences (*p* < 0.05, one-way ANOVA test; n=3). The experiment was repeated three times with similar results. (E, F) Relative expression levels of *SARD1* and *FMO1* in WT and *OX-SNIPER1* plants upon 1 μM nlp20 (E) or 1 μM flg22 (F) treatment. Total RNA was isolated from 12-d-old plate-grown seedlings 4 hours after spraying with 1 μM nlp20 (A) or 1 μM flg22 (B). *ACT1* was used for normalization, and the expression of each gene in the H_2_O (mock)-treated WT plants was set as 1. Error bars represent SD from three different biological replicates. Different letters indicate genotypes with statistical differences (*p* < 0.05, one-way ANOVA test; n=3). The experiment was repeated twice with similar results. (G, H) Levels of free SA in WT and OX-*SNIPER1* plants upon 1 μM nlp20 (G) or 1 μM flg22 (H) treatment. 25-d-old soil-grown plants samples were collected 24 hours after treatment with nlp20, and 9 hours after treatment with flg22. H_2_O served as mock treatment. Different letters indicate genotypes with statistical differences (*p* < 0.05, one-way ANOVA test; n=3). The experiment was repeated three times with similar results. (I) Growth of *Hpa* Noco2 on the local leaves of WT and *OX-SNIPER1* plants with or without nlp20 treatment. Three-week-old plants were pretreated with water (H_2_O) or 1 μM nlp20 and sprayed with *Hpa* Noco2 spores (50,000 spores/ml) 24 hours later. Infection was scored at 7 days post inoculation (dpi) by counting the number of conidiophores per infected leaf. A total of 15 plants were scored for each treatment. Disease rating scores are as follows: 0, no conidiophores on the infected leaves; 1, no more than 5 conidiophores on one infected leaf; 2, 6 to 20 conidiophores on one infected leaf; 3, 20 or more conidiophores on one infected leaf; 4, 5 or more conidiophores on two infected leaves; 5, 20 or more conidiophores on two infected leaves. This experiment was repeated twice with similar results. (J) Growth of *Hpa* Noco2 on the distal leaves of WT and OX-*SNIPER1* plants in an SAR assay. 15 plants were used for each treatment. Disease symptoms were scored 7 dpi by counting the number of conidiophores on the distal leaves. Disease ratings: 0, no conidiophores on plants; 1, one leaf is infected with no more than five conidiophores; 2, one leaf is infected with more than five conidiophores; 3, two leaves are infected but with no more than five conidiophores on each infected leaf; 4, two leaves are infected with more than five conidiophores on each infected leaf; 5, more than two leaves are infected with more than five conidiophores. The experiment was repeated twice with similar results.

### Immune responses induced by nlp20 or flg22 are attenuated in *SNIPER1* overexpression lines

The PTI defects of *SNIPER1* overexpression lines prompted us to test whether *SNIPER1* overexpression affects immune responses induced by the specific PAMP elicitors nlp20 and flg22. To our surprise, the induction of *SARD1* and *FMO1* by nlp20 or flg22 treatment was greatly reduced in the *SNIPER1* overexpression lines compared with WT (Figure 1E and 1F). In addition, SA and SAG levels after nlp20 and flg22 treatment were also significantly lower in the *SNIPER1* overexpression lines than in WT (Figure 1G and 1H, S1B and S1C).

Treatment of nlp20 in local tissues can induce disease resistance in local and distal tissue upon subsequent infection^8^. To determine whether overexpression of *SNIPER1* affects nlp20-induced immunity, we first infiltrated two local leaves with 1 μM nlp20 and sprayed the whole plants with spores of virulent oomycete *Hyaloperonospora arabidopsidis* (*Hpa*) Noco2 one day later, nlp20 treatment induced strong resistance against *Hpa* Noco2 in both the local and distal leaves of WT plants (Figure 1I and 1J). However, the nlp20-induced local as well as systemic resistance to *Hpa* Noco2 were largely impaired in the *SNIPER1* overexpression lines. Together, these data revealed that a general reduction of sNLRs levels due to *SNIPER1* overexpression results in compromised nlp20 and flg22-induced immune responses.

### TNL Signaling components are required for defense against *Pto* DC3000 *hrcC*

As SNIPER1 has been shown to target several TNLs for ubiquitination and degradation^35^, we further tested whether activation of TNL signaling is required for PTI. *Pto* DC3000 *hrcC*–induced defense responses were examined in TNL signaling mutants including *eds1-24, pad4-1, sag101-1, adr1 adr1-L1 adr1-L2* (*adr1 triple*) and *nrg1a nrg1b nrg1c* (*nrg1 triple*) mutants. The induction of *SARD1* by *Pto* DC3000 *hrcC* was almost completely blocked in *eds1-24, pad4-1* and *adr1 triple* mutant plants, while no change in induction was observed in the *sag101-1* and *nrg1 triple* mutants compared with the WT (Figure S2A). Similarly, the induction of *FMO1* is greatly reduced in *eds1-24, pad4-1* and *adr1 triple*, but hardly affected in *sag101-1* and *nrg1 triple* mutant plants (Figure S2B).

We further measured SA accumulation following *Pto* DC3000 *hrcC* infection in these TNL signaling mutants. Both SA and SAG levels in *eds1-24, pad4-1* and *adr1 triple* after *Pto* DC3000 *hrcC* treatment were much lower than in the WT (Figure 2A, S2C). This is consistent with the known contributions of EDS1, PAD4 and the ADR1s to pathogen-induced SA biosynthesis. To our surprise, the SA levels after *Pto* DC3000 *hrcC* treatment were also significantly reduced in *sag101-1* and *nrg1 triple* mutant plants, although the reduction was not as dramatic as in *eds1-24*, *pad4-1* and *adr1 triple*. Consistent with the difference in SA levels, growth of *Pto* DC3000 *hrcC* was significantly higher in *sag101-1* and *nrg1 triple* than in WT, and further increased in *eds1-24, pad4-1* and *adr1 triple* leaves (Figure 2B). Taken together, PTI responses are significantly attenuated in TNL signaling mutants, indicating a key contribution of TIR signaling to PTI.

**Figure 2.**
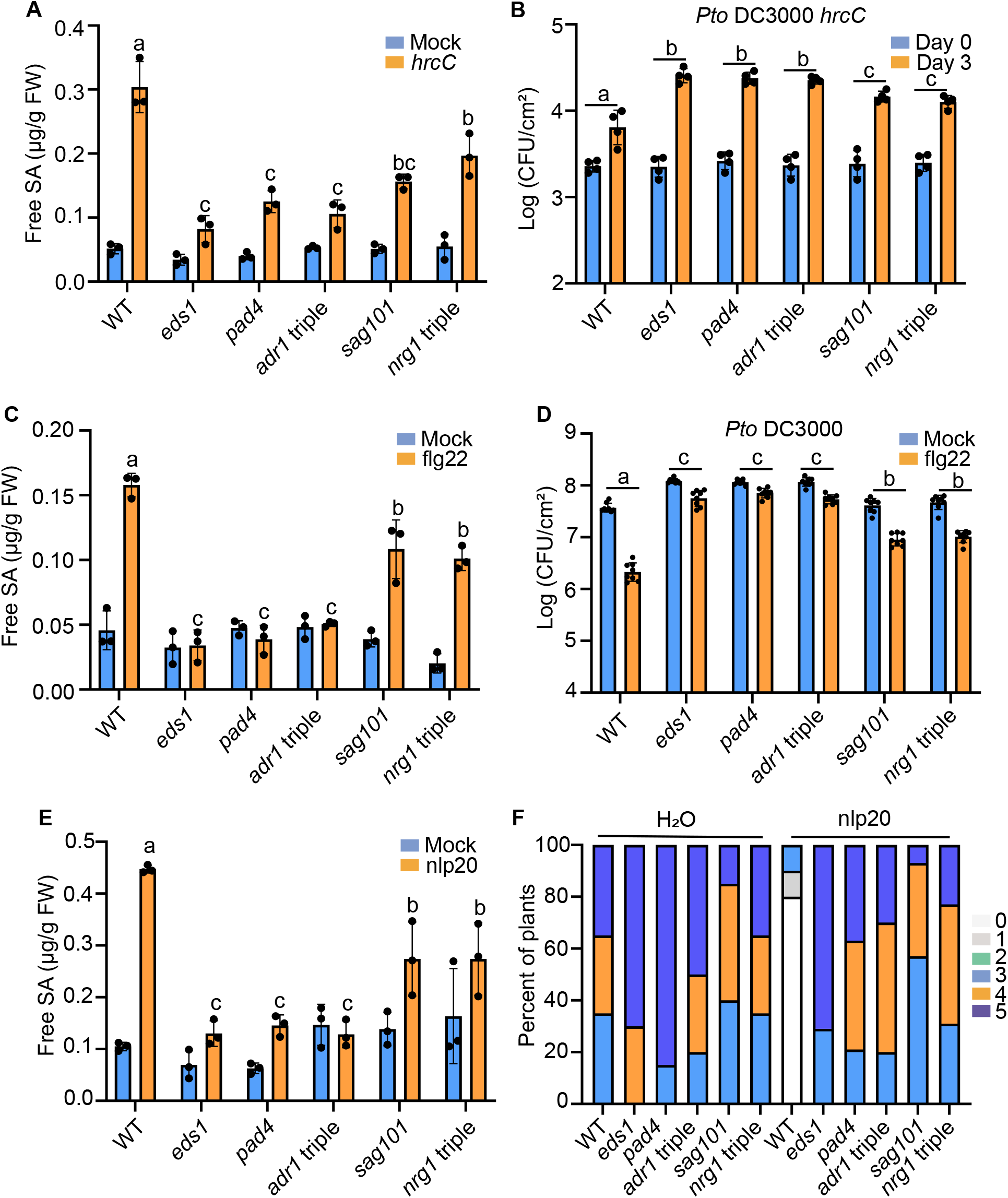
Contributions of TIR signaling components to *Pto* DC3000 *hrcC*, flg22 and nlp20-induced immunity. (A) Levels of free SA in four-week-old soil-grown WT, *eds1-24, pad4-1, sag101-1, adr1 triple* and *nrg1 triple* mutants 12 hours after treatment with 10 mM MgCl2 or *Pto* DC3000 *hrcC* (OD_600_=0.05) for 12 hours. Different letters indicate genotypes with statistical differences (*p* < 0.05, one-way ANOVA test; n=3). The experiment was repeated three times with similar results. (B) Growth of *Pto*DC3000 *hrcC* in 25-day-old soil-grown plants of the indicated genotypes. Leaf discs were collected 0 days (Day 0) or 3 days (Day 3) after *Pto* DC3000 *hrcC* (OD_600_=0.002) infiltration. Error bars represent SD from four biological replicates. The growth of *Pto* DC3000 *hrcC* in different genotypes was compared using two-way ANOVA test, and different letters indicate genotypes with statistical differences (*p* < 0.05, *n* = 4). The experiment was repeated twice with similar results. (C) Levels of free SA in four-week-old soil-grown plants of the indicated genotypes after treatment with water or 1 μM flg22 for 9 hours. Different letters indicate genotypes with statistical differences (*p* < 0.05, one-way ANOVA test; n=3). The experiment was repeated three times with similar results. (D) Growth of *Pto* DC3000 in the leaves of four-week-old WT, *eds1-24, pad4-1, sag101-1*, *adr1 triple* and *nrg1 triple* mutant plants after treatment with water or 1 μM flg22. 24 hours later, the same treated leaves were infiltrated with *Pto* DC3000. Samples were taken 3 days after *Pto* DC3000 inoculation. Error bars represent SD from six biological replicates. The flg22-induced protection among different genotypes was compared using two-way ANOVA test, and different letters indicate genotypes with statistical differences (*p* < 0.05, *n*= 6). The experiment was repeated twice with similar results. (E) Levels of free SA in four-week-old soil-grown plants of the indicated genotypes after treatment with water (mock) or 1 μM nlp20 for 24h. Different letters indicate genotypes with statistical differences (*p* < 0.05, one-way ANOVA test; n=3). The experiment was repeated three times with similar results. (F) Growth of *Hpa* Noco2 on the local leaves of the indicated plants. Three-week-old soil-grown plants were pretreated with water or 1 μM nlp20 and sprayed with *Hpa* Noco2 spores (50,000 spores/ml) 24 hours later. Disease ratings are as described in Fig 1. The experiment was repeated twice with similar results.

### flg22-induced immune responses are attenuated in TNL signaling mutants

We then tested PTI responses induced by specific elicitors in the TNL signaling mutants. To determine whether flg22-induced defense responses require TIR signaling, we first compared *SARD1* and *FMO1* induction by flg22 treatment. The expression levels of both *SARD1* and *FMO1* after flg22 treatment were considerably lower in *eds1-24, pad4-1* and *adr1 triple* mutant plants (Fig S2D and S2E). In contrast, the expression of *SARD1* was not affected whereas *FMO1* induction was modestly reduced in *sag101-1* and *nrg1 triple*. We also compared the accumulation of SA and SAG after flg22 treatment in WT and the TNL signaling mutants. The SA and SAG levels in *eds1-24, pad4-1* and *adr1 triple* plants treated with flg22 were much lower than in the WT (Figure 2C, S2F). Interestingly, the SA levels in *sag101-1* and *nrg1 triple* were also significantly lower compared to the WT, but the difference is not as dramatic as in *eds1-24, pad4-1* and *adr1 triple* plants. Consistent with the reduced SA levels, flg22-induced resistance against *Pto* DC3000 was also compromised in *eds1-24, pad4-1* and *adr1 triple*, as well as in *sag101-1* and *nrg1 triple* mutant plants, although to a lesser extent (Figure 2D). Taken together, activation of TIR signaling contributes to flg22-induced immune responses.

### nlp20-induced immunity is lost in TNL signaling mutants

To determine whether TIR signal components are required for nlp20-induced immune responses, we first examined nlp20-induced *SARD1* and *FMO1* expression in *eds1-24*, *pad4-1, sag101-1, adr1 triple* and *nrg1 triple* mutants. The induction of *SARD1* and *FMO1* by nlp20 was dramatically reduced in *eds1-24, pad4-1* and *adr1 triple* mutant plants, but hardly affected in *sag101-1* and *nrg1 triple* (Figure S2G and S2H). We then measured nlp20-induced SA accumulation in these TNL signaling mutants. The SA and SAG levels after nlp20 treatment were much lower in *eds1-24, pad4-1* and *adr1 triple* plants, and moderately lower in *sag101-1* and *nrg1 triple* than in WT (Figure 2E and S2I). Consistent with the reduced SA accumulation, nlp20-induced resistance against *Hpa* Noco2 in both local and distal tissue was attenuated in *sag101-1* and *nrg1 triple* and almost completely blocked in *eds1-24, pad4-1* and *adr1 triple* (Figure 2F and S3). Taken together, activation of TIR signaling is required for nlp20-induced immunity.

### Overexpression of genes encoding TIR domain-containing proteins activates defense responses

To understand how TIR signaling is activated during PTI, we analyzed the expression of genes encoding TIR domain-containing proteins (*TIR* genes) in response to nlp20 or flg22 using previously reported RNA-sequencing datasets^36,37^. 26 *TIR* genes were found to be significantly induced 1 h after nlp20 treatment (Table S1). 14 *TIR* genes were considerably induced 6 h after applying nlp20 (Table S2). With flg22 treatment, 46 TIR genes were significantly up-regulated within 30 min (Table S3). These findings suggest that a large number of TIR genes are rapidly induced upon PTI activation.

To test whether up-regulation of *TIR* genes can activate defense responses, we transiently expressed three *TIR* genes induced by both nlp20 and flg22 in *Nicotiana benthamiana*. Among them, *AT4G11170* and *AT3G04220* encode full-length TNLs and *AT2G32140* encodes a protein with only the TIR domain. Overexpression of all three *TIR* genes in *N. benthamiana* leads to activation of cell death around 48 hours after *Agrobacteria* infiltration (Figure 3A). To determine whether overexpression of these three *TIR* genes activates SA biosynthesis, we measured SA levels in samples collected 24 hours and 36 hours after infiltration of the *Agrobacteria* strains, when no macroscopic cell death was visible. In agreement, overexpression of these *TIR* genes in *N. benthamiana* indeed resulted in dramatic increase in SA and SAG levels in *N. benthamiana* (Figure 3B, S4).

**Figure 3.**
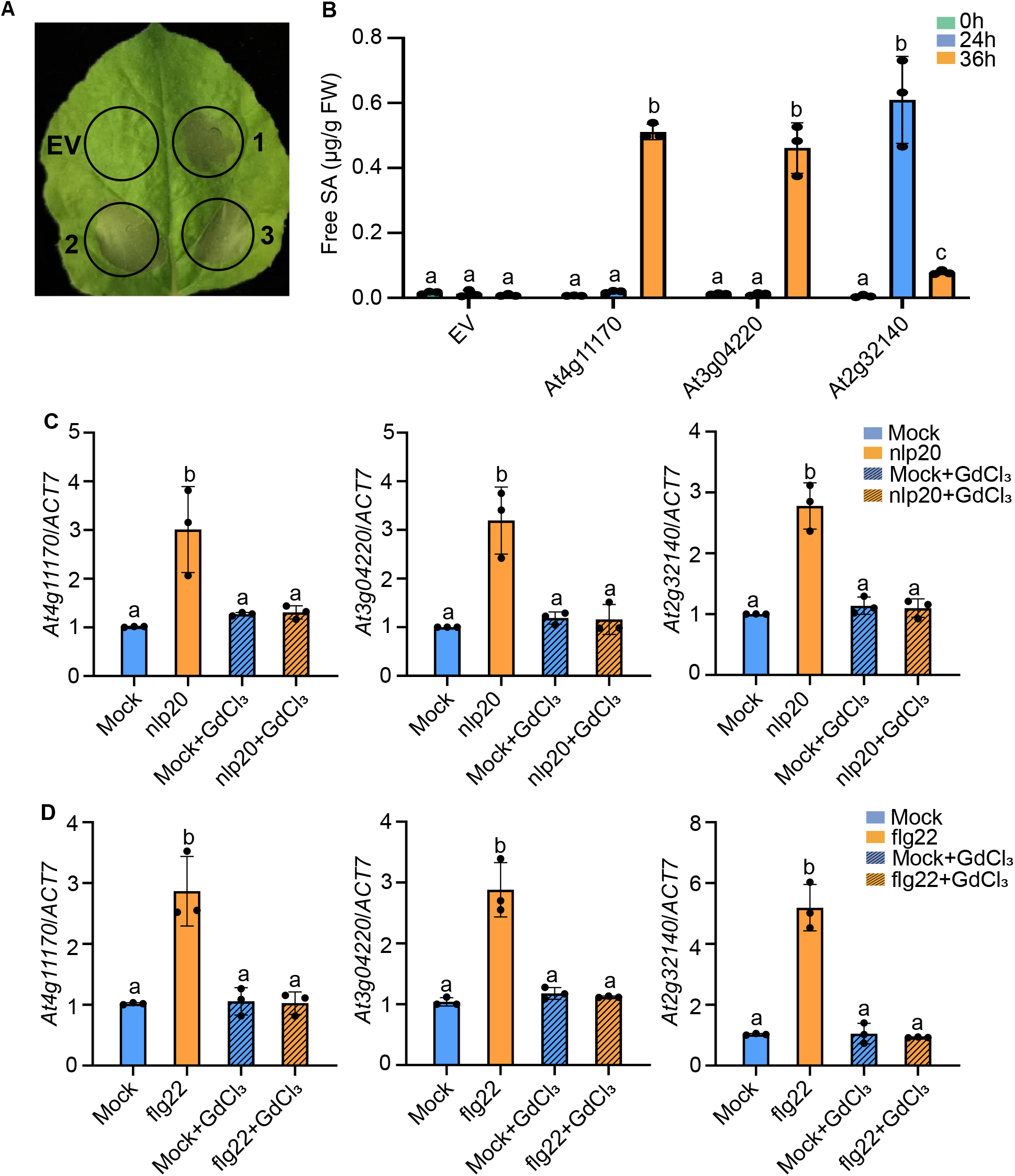
Overexpression of *TIR* genes activates SA biosynthesis in *N. benthamiana*. (A) Hypersensitive response (HR) in the *N. benthamiana* leaves expressing the TIR genes *At4g11170, At3g04220* or *At2g32140* through *Agrobacterium tumefaciens* GV3101 (OD_600_=0.4) infiltration. Photographs were taken 3 days after *Agrobacterium* infiltration. The empty vector (EV) treatment serves as control; 1, *At4g11170*; 2, *At3g04220*; 3, *At2g32140*. (B) Levels of free SA in *N. benthamiana* leaves after infiltration of *Agrobacterium* (OD_600_=0.4) carrying the *TIR* genes from (A). Samples were collected 24 and 36 hours post infiltration, before HR was visible. Error bars represent SD from three biological replicates. Different letters indicate different time course with statistical differences (*p* < 0.05, one-way ANOVA test; n=3). (C, D) Induction of the indicated *TIR* genes by nlp20 (C) or flg22 (D). Ten-day--old plate-grown WT plants were transplanted to water 1 day before for recovery and then pretreated with water (Mock) or 100 μM GdCl_3_ for 1 hour. Samples were collected 1 hour after supplying with 1 μM nlp20 or 1 μM flg22. qPCR was used to examine the genes expression level. *ACT7* was used for normalization. Error bars represent SD from three different biological replicates. Different letters indicate different treatment with statistical differences (*p* < 0.05, one-way ANOVA test; n=3) All the experiments were repeated twice with similar results.

Since Ca^2+^ influx is one of the earliest events during PTI, we further examined whether it is involved in activation of the *TIR* genes. To determine whether Ca^2+^ influx is required for nlp20-induced up-regulation of the three *TIR* genes, we pretreated *Arabidopsis* seedling with GdCl_3_ to block the Ca^2+^ channels prior to nlp20 treatment. Consistent with the RNA-sequencing datasets, treatment with nlp20 or flg22 alone leads to rapid induction of the three *TIR* genes. However, this induction was completely blocked by GdCl_3_ (Figure 3C-D), suggesting that activation of Ca^2+^ signaling is required for the induction of *TIR* genes.

### nlp20-induced immunity requires the RLCKs PCRK1/2 and PBL19/20

RLCKs PCRK1/2 and PBL19/20 were known to function downstream of PRR receptor kinases such as FLS2 and CERK1^38–40^. To determine whether they are required for nlp20-induced immunity, we compared growth of *Hpa* Noco2 on WT, *pcrk1/2, pcrk1/2 pbl19* and *pcrk1/2 pbl19/20* quadruple mutant plants after treatment with nlp20. nlp20-induced local and systemic resistance against *Hpa* Noco2 was compromised in *pcrk1/2* and *pcrk1/2pbl19*, and almost completely blocked in *pcrk1/2 pbl19/20* (Figure 4A, S5A). Similarly, nlp20-induced resistance against *Pto* DC3000 was also severely compromised in the *pcrk1/2pbl19/20* quadruple mutant (Figure S5B). In addition, the increased SA and SAG levels after nlp20 treatment were significantly reduced in *pcrk1/2pbl19/20* than in the WT (Figure 4B and S5C). Further RT-qPCR analysis showed that nlp20-induced *SARD1* and *FMO1* expression was dramatically reduced in *pcrk1/2 pbl19/20* mutant plants (Figure S5D and S5E). Moreover, induction of the three above-mentioned TIR genes was blocked in *pcrk1/2 pbl19/20* (Figure 4C). Together these data indicate that PCRK1/2 and PBL19/20 are required for nlp20-induced immunity and they act upstream of the early induction of *TIR* genes.

**Figure 4.**
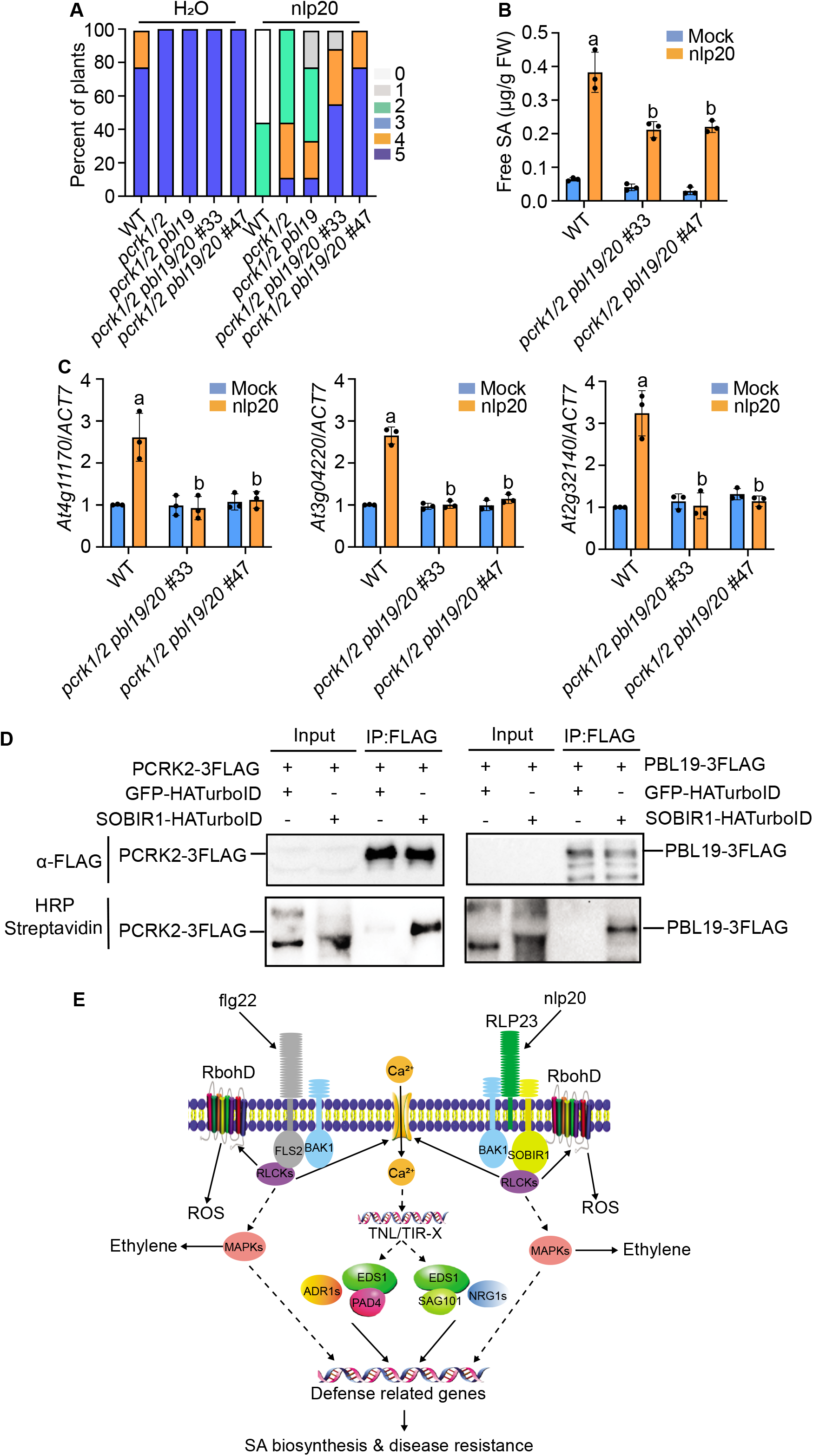
PCRK1/2 and PBL19/20 are required for nlp20-induced immunity. (A) Growth of *Hpa* Noco2 on the distal leaves of WT, *pcrk1/2, pcrk1/2 pbl19, pcrk1/2 pbl19/20* #33 and *pcrk1/2pbl19/20* #47 quadruple mutant plants. 21-d-old soil-grown plants were treated with 1 μM nlp20 and sprayed *Hpa* Noco2 spores (50,000 spores/ml) 24 hours later. The detail method was described in Fig 2F. (B) Levels of free SA in four-week-old soil-grown plants of the indicated genotypes treated with water or 1 μM nlp20. Samples were collected 24 hours post elicitor treatment. Different letters indicate genotypes with statistical differences (*p* < 0.05, one-way ANOVA test; n=3). (C) The induction of *At4g11170, At3g04220* and *At2g32140 (TIR* genes) in the indicated genotypes. Total RNA was isolated from seedlings of 10-d-old plate-grown plants 1 h after treatment with 1 μM nlp20. *ACT7* was used for normalization, and the expression of each gene in mock-treated WT was set as 1. Error bars represent SD from three different biological replicates. Different letters indicate genotypes with statistical differences (*p* < 0.05, one-way ANOVA test; n=3). (D) Immunoprecipitation and biotinylation of *PCRK2/PBL19-3FLAG* by *SOBIR1-HATurboID* in *N. benthamiana. Agrobacterium* carrying the indicated constructs were infiltrated into *N. benthamiana* leaves for protein expression. Immunoprecipitation was carried out with anti-FLAG beads. The 3FLAG-tagged proteins were detected using an anti-FLAG antibody. The biotinylated proteins were detected using HRP-Streptavidin. The experiments in (A-D) were repeated twice with similar results. (E) A working model for the contribution of TIR signaling in PTI. Upon perception of pathogen elicitors such as flg22 and nlp20, PRRs activate early immune responses such as ROS production, MAPK activation and Ca^2+^ influx through RLCKs. Elevated cytosolic Ca^2+^ levels induce the expression of a large number of *TIR* genes, leading to activation of downstream defense pathways through the EDS1/PAD4/ADR1s and EDS1/SAG101/NRG1s signaling modules. Activation of TIR signaling further induces downstream defense gene expression, resulting in increased SA biosynthesis and enhanced resistance to pathogens. In parallel, activation of MAPKs promotes the biosynthesis of ethylene.

Since SOBIR1 works together with the nlp20 receptor RLP23 in nlp20-activated PTI signal transduction^8^, one question is whether PCRK1/2 and PBL19/20 function immediately downstream of SOBIR1. Therefore, we tested whether SOBIR1 directly interacts with PCRK2 and PBL19 using TurboID, a highly efficient proximity labeling method for detecting protein-protein interactions^34,41^. The SOBIR1-HA-TurboID fusion protein was co-expressed with 3×FLAG-tagged PCRK2 or PBL19 in *N. benthamiana*. After biotin treatment, the 3 ×FLAG-tagged PCRK2 or PBL19 proteins were immunoprecipitated to examine their biotinylation using Streptavidin-HRP. Both PCRK2 and PBL19 were biotinylated by SOBIR1-HA-TurboID, suggesting that SOBIR1 directly interacts with PCRK2 and PBL19 (Figure 4D). These data agree with the general notion of RLCKs acting immediately downstream of PRRs. We also tested whether SOBIR1 interacts with EDS1/PAD4/ADR1 using TurboID. However, no biotinylation or co-immunoprecipitation of EDS1, PAD4 or ADR1 by SOBIR1-HA-TurboID was observed (Figure S6).

## Discussion

PTI and ETI have been traditionally studied as separate defense pathways. Recently it was reported that PTI is required for the activation of NLR-mediated ETI^42,43^. Here, we showed that loss of TIR signaling as well as reduced NLR accumulation results in compromised defense responses activated by nlp20, flg22 and *Pto* DC3000 *hrcC*, indicating that activation of TIR signaling is essential for PTI. While the EDS1/PAD4/ADR1s module plays a predominant role, the EDS1/SAG101/NRG1 module also contributes to flg22, nlp20 and *Pto* DC3000 *hrcC*-induced plant immunity.

SA plays crucial roles in resistance against biotrophic pathogens such as *Hpa* Noco2 and *Pto* DC3000. Activation of PTI leads to rapid increase of SA biosynthesis and elevated SA levels^9^. In the *SNIPER1* overexpression lines, SA levels following nlp20 treatment and *Pto* DC3000 *hrcC* infection are much lower compared to the WT plants. Similarly, nlp20 and *Pto* DC3000 *hrcC*-induced SA accumulation is greatly reduced in *eds1-24, pad4-1* and *adr1 triple* mutant plants. In *nrg1 triple* and *sag101-1* mutant plants, SA levels after nlp20 treatment and *Pto* DC3000 *hrcC* infection are also significantly lower than in the wild type. These findings suggest that both the EDS1/PAD4/ADR1s and EDS1/SAG101/NRG1s modules downstream of TIR-type receptors contribute to the up-regulation of SA levels, which plays important roles in defense against pathogens.

In flg22-treated *adr1 triple* mutant plants, SA levels are significantly reduced compared to WT^22^. However, other early PTI responses such as ROS production, MAPK activation and callose deposition induced by flg22 and elf18 are not affected in the *adr1 triple* mutant, suggesting that TIR signaling mediated by ADR1s is not required for the induction of all the very early PTI responses^22^. In flg22-treated *eds1-24, pad4-1, adr1 triple, sag101-1* and *nrg1 triple* mutant plants, flg22-induced resistance against *Pto* DC3000 is significantly reduced but not completely blocked. The loss of TIR signaling is likely compensated by other defense pathways downstream of FLS2, as combining *pad4-1* with the JA biosynthesis mutant *dde2-2*, the ethylene response mutant *ein2-1* (*ethylene insensitive 2-1*) and the SA biosynthesis mutant *sid2-2* (*salicylic acid induction deficient 2-2*) leads to a complete loss of flg22-induced protection against *Pto* DC3000^44^.

In mock-treated *adr1 triple, eds1-24* and *pad4-1* mutants, growth of *Pto* DC3000 is much higher than in wild type plants. *adr1 triple, eds1-24* and *pad4-1* mutant plants are also more susceptible to *Hpa* Noco2. The general enhanced susceptibility of these mutants to virulent pathogens could be due to compromised PTI caused by loss of the reinforcement of defense responses through activation of TIR signaling.

*SARD1* encodes as a master transcription factor regulating the expression of a large number of defense regulators as well as genes involved in SA biosynthesis^45^. Overexpression of *SARD1* leads to increased SA levels and enhanced disease resistance^29^. In WT plants, the expression of *SARD1* is rapidly and strongly induced during PTI. Interestingly, the induction of *SARD1* by nlp20, flg22 and *Pto* DC3000 *hrcC* is dramatically reduced in the *SNIPER1* overexpression lines and mutants deficient in TIR signaling such as *adr1 triple, eds1-24* and *pad4-1*, suggesting that the *SARD1* induction during PTI depends on the activation of TIR signaling, which is consistent with the reduced SA levels in the *SNIPER1* overexpression lines and TIR signaling mutants.

A large number of *TIR* genes including *SNC1* and *LAZ5* are rapidly induced after treatment with nlp20 or flg22 (Table S1–S3). Overexpression of *SNC1* and *LAZ5* is known to result in constitutively activated defense responses^46,47^. Similarly, overexpression of the TIR-X protein encoded by *At2g32140* also leads to constitutive activation of EDS1 and PAD4-dependent immune responses^48^. In addition, overexpression of the TIR domains of many TNLs alone is sufficient to activate cell death in *N. benthamiana*^14,15,49^. Recently, many TIR domains have been shown to exhibit NADase activity, likely generating signal molecule(s) activating EDS1-dependent defense responses^14,15^. Induction of the TIR genes during PTI could lead to increased production of these defense signal molecules and activation of downstream TIR signaling pathways and increased SA biosynthesis (Figure 4E). In support of this, transient overexpression of *AT2G32140* and the TNL genes *At4G11170* and *AT3G04220*, which are up-regulated during PTI, resulted in dramatic increase in SA accumulation.

The RLCKs PCRK2 and PBL19 were found to directly interact with SOBIR1. Together with two other closely related RCLKs PCRK1 and PBL20, they are required for nlp20-induced resistance against *Hpa* Noco2. In *pcrk1/2pbl19/20* quadruple mutant plants, nlp20-induced *SARD1* expression and SA production was blocked. In addition, the induction of several *TIR* genes by nlp20 is also abolished in the *pcrk1/2 pbl19/20* mutant. These findings suggest that PCRK1/2 and PBL19/20 function downstream of the RLP23/SOBIR1 receptor complex to activate the expression of nlp20-responsive *TIR* genes, leading to activation of TIR signaling and SA biosynthesis (Fig 4E). Interestingly, three other RLCKs, PBL30/31/32, were recently reported to be required for nlp20-induced ROS and ethylene production and resistance to *Pto* DC3000^50^. It is possible that different RLCKs activate different downstream components of the RLP receptor complex, leading to branching of the downstream defense pathways.

In summary, in addition to effector recognition, some TNLs and TIR-X proteins also play important roles in amplifying PTI responses (Figure 4E). Activation of TIR signaling during PTI is most likely through the induction of *TIR* genes. How the expression of the *TIR* genes is activated is currently unclear. As the induction of several *TIR* genes by nlp20 is blocked by the Ca^2+^ channel blocker GdCl_3_, elevation in cytosolic Ca^2+^ levels caused by activation of Ca^2+^ channel(s) during PTI may play a crucial role in *TIR* gene induction (Figure 4E). The identities of the transcription factors involved in up-regulation of the *TIR* genes and the mechanism of how Ca^2+^ may affect their activities remains to be determined. Whether PCRK1/2 and PBL19/20 are involved in activation of the Ca^2+^ channel(s) also needs to be determined in the future.

## Materials and methods

### Plasmid constructs

To generate the CRISPR/Cas9 construct for genome editing of *EDS1A/B* and *PBL20*, genomic sequences of *EDS1* and *PBL20* were subjected to CRISPRscan (http://www.crisprscan.org/?page=sequence) to identify the target sequences. The selected sequences were evaluated with Cas-OFFinder (http://www.rgenome.net/cas-offinder/). The target sequences used for editing *EDS1A/B* was 5’-CTAACCGAGCGCTATCACA(AGG)-3’ and 5’-CGGAGAATACATCTCCCTT(TGG)-3’, for PBL20 was 5’-CCAAAATCCAGAGGAAATA(TGG)-3’ and 5’-CAATAAGTATCCAATTGCTA(TGG)-3’. CRISPR constructs were generated in the *pHEE401E* vector using a previously described CRISPR-Cas9 gene editing system^51^.

For coimmunoprecipitation, *PCRK2, PBL19* and *ADR1* were amplified by primers PCRK2-Kpn1-F and PCRK2-Spe1-R, PBL19-Kpn1-F and PBL19-BamH1-R, or ADR1-KpnI-F and ADR1-SalI-R, then cloned into pBASTA-35S-3FLAG vector. The *SOBIR1* fragment was cut from pBASTA-35S-SOBIR1-3FLAG plasmid^52^, then sub-cloned into pBASTA-35S-2HA-Turbo vector. *EDS1* and *PAD4* were first amplified by primers EDS1-KpnI-F and EDS1-XbaI-R or PAD4-Kpn1-F and PAD4-BamHI-R, and cloned into pCambia1305-FLAG-ZZ vector^53^.

### Plant materials and growth conditions

The *pad4-1, sag101-1, adr1 triple, nrg1 triple*, and *pcrk1 pcrk2 (pcrk1/2)* double mutants were previously described^20,22,38,54,55^. The *eds1-24* deletion line was generated by transformation of a *EDS1* CRISPR construct into WT Col-0 plants. Deletion and presence primers were used to detect the presence and homozygosity of the deletion (Supplemental Table 4). The *eds1-24* line is a Cas9 transgene-free line homozygous for a 2636bp deletion, causing truncations of both *EDS1A* (*AT3G48090*) and *EDS1B* (*AT3G48080*).

The *pcrk1 pcrk2 pbl19* (*pcrk1/2 pbl19*) triple mutant was generated by crossing *pcrk1 pcrk2* with *pbl19-2* (Salk_065136C). The *pcrk1 pcrk2 pbl19 pbl20* (*pcrk1/2pbl19/20*) quadruple mutants were generated by transforming the CRISPR/Cas9 construct targeting *PBL20* into the *pcrk1 pcrk2 pbl19* triple mutant background. Both *pcrk1 pcrk2 pbl19 pbl20* #33 and #47 lines carry a large 1.5 kb deletion in *PBL20*. All the mutants are in the Col-0 background. The transgenic OX-*SNIPER1* #4, OX-*SNIPER1* #5 lines were generated previously in a reverse genetics screen for plant immunity related E3 ligases^20^.

Plants were grown in growth rooms with a temperature of 23°C under long day (16h light/8h dark) or short day (12h light/12h dark) condition at approximately 100 μmol m^-2^ s^-1^ light intensity. For *Agrobacterium* mediated transformation, the *Arabidopsis* seeds were directly sown on soil and grown for around 5 weeks prior to floral-dip transformation. For RNA isolation, the *Arabidopsis* seeds were sterilized in 15% (v/v) bleach and germinated on plates with ½ Murashige and Skoog (MS) with vitamins (PlantMedia) and 1% (w/v) sucrose.

### RNA extraction and gene expression

For analyzing nlp20/flg22 induced gene expression, total RNA was extracted from 12-day-old plate-grown seedlings 4 hours after spraying 1 μM flg22 or 1μMnlp20. To test pathogen-induced gene expression, leaves of four-week-old plants grown under short day conditions were infiltrated with *Pto* DC3000 *hrcC* at a dose of OD_600_=0.05 and collected after 12hours. RNA was extracted using the EZ-10 Spin Column Plant RNA Mini-Preps Kit (Bio Basic, Canada) according to the manufacturer’s instructions. 1 μg RNA was used for cDNA synthesis by Oligo(dT)-primed reverse transcription using the OneScript Reverse Transcriptase kit (ABM, Canada). Real-time quantitative PCR was performed to analyze the gene expression levels, using the SYBR Premix Ex Taq II kit (TAKARA). The primers used for qPCR were reported previously^45^. *ACTIN1* was used as an internal control. For nlp20-induced *TIR* gene expression analysis, 10-day-old seedlings grown on ½ MS medium were transferred to ½ MS liquid medium. After 24-hour recovery, nlp20 was supplied to a 1 μM final concentration. Two to three individual plants treated with nlp20 were harvested 1 hour later as one sample. *ACTIN7* was used as an internal control. The primers used are listed in Supplemental Table 4.

### Measurement of Salicylic acid

The procedure for SA extraction and measurement was reported previously^56^. In brief, for each sample, about 100mg leaf tissue was collected from 25-d-old soil-grown plants and grounded in liquid nitrogen. Each genotype contains four biological replicates, with each sample collected from three different plants. For every sample, 0.6 ml of 90% methanol was added, and the sample was vortexed 20s and sonicated for 20 min to release SA. Samples were then centrifuged at 12000×g for 10 min. The supernatant was collected and another 0.5 ml of 100% methanol was added to the pellet for a second round of extraction. The supernatant from both extractions were combined and dried by vacuum. Then 0.5 ml 5% (w/v) trichloroacetic acid was added to the dry samples, vortexed and sonicated for 5 min, and then centrifuged at 12000×g for 15 min. The supernatant was collected and then extracted three times with 0.5 ml extraction buffer (ethylacetate acid/ cyclopentane/ isoporopanal: 100/99/1 by volume).

Each time, after centrifugation at 12,000×g for 10 min, the organic phase was collected and combined to a new tube and dried by vacuum afterwards. The final dried sample was resuspended in 200 μl mobile phase (0.2M KAc, 0.5mM EDTA PH=5) by vortexing and sonicating for 5 min. After spinning at 12,000×g for 5 min, the supernatant was kept and measured by high-performance liquid chromatography (HPLC) to measure the amount of SA as compared with a standard.

### Pathogen infection assay

For *Pto* DC3000 *hrcC* bacterial growth assays, two fully extended leaves of four-week-old plants grown under short-day conditions were infiltrated with *Pto* DC3000 *hrcC* at a dose of OD_600_ = 0.002. Samples were collected at 0 day and 3 days after infiltration. One sample contained two leaf discs from a single plant, and a single leaf disc was collected from each infected leaf. The samples were ground, diluted in 10 mM MgCl2, and plated on Lysogeny broth (LB) agar plates containing 25 μg ml^-1^ Rifampicin. After incubation at 28C’ for 36 h, colonies were counted from selected dilutions and the colony numbers were used to calculate the colony forming units. For flg22/nlp20-induced pathogen resistance, leaves of four-week-old plants were infiltrated with 1 μM flg22/nlp20 or H_2_O as control. After 24h, the same treated leaves were infiltrated with *Pto* DC3000 at a dose of OD_600_=0.001. After 3 days, samples were collected and analyzed as above.

To analyze nlp20-induced SAR, leaves of three-week-old plants were first infiltrated with 1 μM nlp20. After 24 h, plants were sprayed with *Hpa* Noco2 spore suspension at a concentration of 50,000 per ml water. Then plants were covered with a clean dome and grown at 18°C under short-day conditions in a growth chamber. After 7 days, the *Hpa* Noco2 sporulation was scored as previously described^57^.

### TurboID-based proximity labeling in *N. benthamiana*, immunoprecipitation and western blot analysis

TurboID-based proximity labeling assay was performed as described previously^20^. In brief, *N. benthamiana* leaves were infiltrated with *Agrobacterium* containing *HA-TurboID* and *ZZ-TEV-FLAG* or *3xFLAG* tagged constructs. At 48 hpi, biotin was infiltrated, and the plants were incubated at room temperature for 2 hours to allow biotin labeling. About 2.0 g *N. benthamiana* leaves expressing the indicated proteins were harvested at 50 hpi and ground into powder with liquid nitrogen. Extraction buffer containing 25 mM Tris-HCl pH 7.5, 150 mM NaCl, 1mM EDTA, 0.3% Nonidet P-40, 10% Glycerol, 1 mM PMSF, 1× Protease Inhibitor Cocktail (Roche; Cat. #11873580001), and 10 mM DTT. The FLAG-tagged PCRK2 and PBL19 proteins were immunoprecipitated using 15 μl M2 beads (Sigma; Cat. #A2220). Biotinylation was detected with Streptavidin-HRP (Abcam Cat. # ab7403). The anti-HA antibody was from Roche (Cat. #11867423001). The anti-FLAG antibody was from Sigma (Cat. #F1804).

### Bioinformatic analysis for TIR-containing gene induction

TIR genes induced 30 minutes after flg22 treatment were subset from previously published data^58^. For nlp20-induced genes, previously published raw RNA-seq reads were retrieved (GSE133053)^36^. BBDuk (https://sourceforge.net/projects/bbmap/) was used to trim adapters. A decoy-aware reference transcriptome was generated using a high-quality *Arabidopsis* reference transcriptome, AtRTDv2_QUASI_19April2016.fa^59^, and an *Arabidopsis* whole genome sequence (Ensembl Plants version 47) as a decoy. Salmon v1.2.1^60^ was used to build an index and quantify transcript expression against the reference transcriptome using default parameters. Transcript-level expression (TPM values) were imported to R and summarized to gene-level expression using tximport v1.16.1^61^. DESeq2 v1.28.1^62^ was used to determine differentially-expressed genes (padj < 0.1). Genes were annotated using biomaRt v2.44.1^63^, and genes containing TIR domains (IPR035897, IPR000157, IPR041340, IPR017279) were subset.

## Author Contributions

HT, SC and ZW carried out the majority of the experiments. KA generated *eds1-24*, and extracted the up-regulated *TIR* genes from RNA-Sequencing datasets. WH and YZ helped with HPLC SA analysis. FX made the FLAG-ZZ tagged EDS1 and PAD4 constructs. HY and TS generated the combined RLCK mutants. YZ, SW and XL wrote the manuscript with contributions of all authors.

## Competing financial interests

The authors declare no competing financial interests.

## Acknowledgements

We would like to thank Drs. Jane Parker, Jeff Dangl and Jian-Min Zhou for sharing mutant seeds and pathogen strains. Mr. Lei Tian is thanked for help with the HPLC analysis. Dr. Thorsten Nuernberger is thanked for insightful discussions. This study was financially supported by grants to XL and YZ from the Natural Sciences and Engineering Research Council (NSERC) Discovery program of Canada, NSERC-CREATE-PRoTECT, and the Canadian Foundation for Innovation (CFI), grant to YZ from National Natural Science Fundation of China (31828008), scholarships to HT, SC, WH and ZW from the Chinese Scholarship Council, and scholarships to KA from the Alexander Graham Bell Canada Graduate Scholarship Doctoral Program, and the University of British Columbia Four-year fellowship program.

## Supplementary tables

**Table S1: TIR genes induced by flg22 30 min after treatment.**

**Table S2: TIR genes induced by nlp20 1h after treatment.**

**Table S3: TIR genes induced by nlp20 6h after treatment.**

**Table S4: Sequence of primers used in this study.**

## Supplementary figure legends

**Figure S1.**
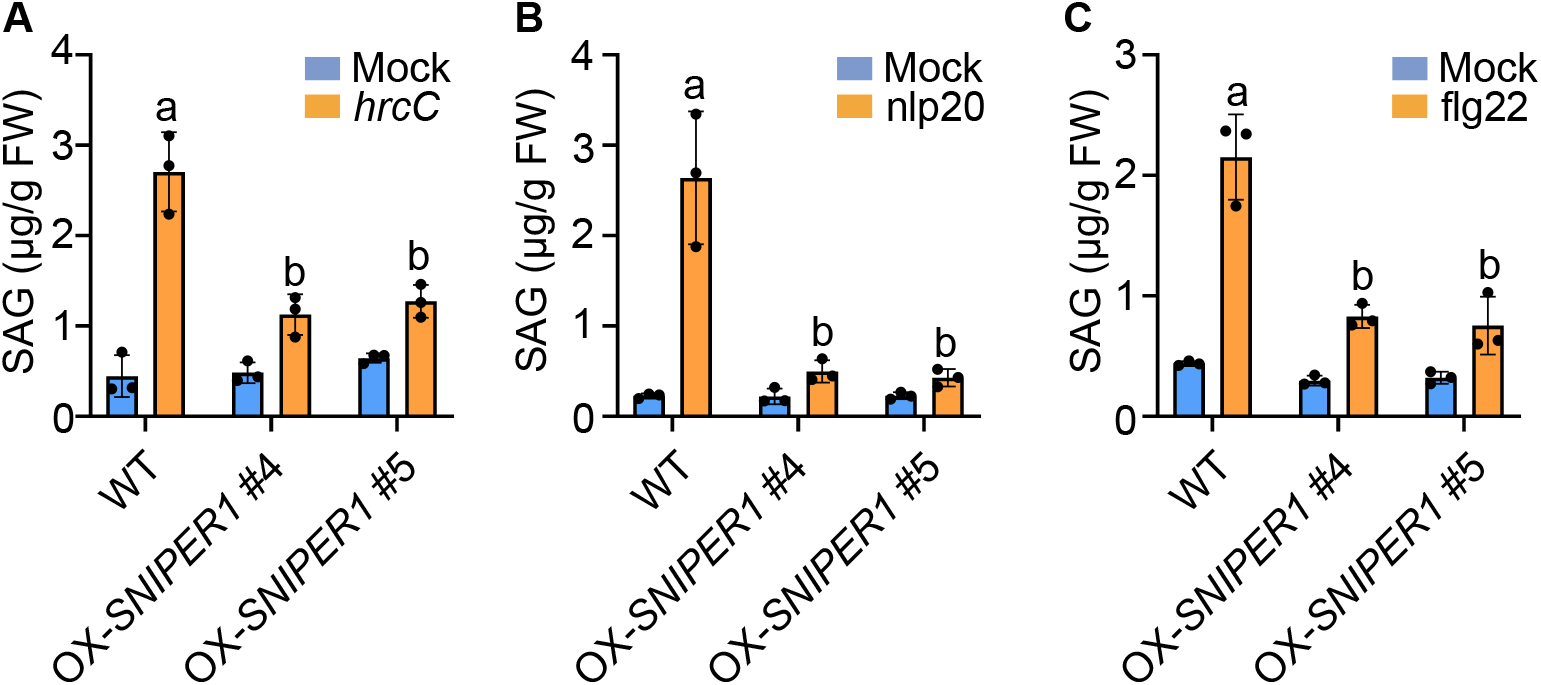
Levels of glucose-conjugated SA (SAG) in WT and *OX-SNIPER1* plants. (A) SAG levels in four-week-old soil-grown WT and OX-*SNIPER1* plants treated with 10 mM MgCl2 (mock) or *Pto* DC3000 *hrcC*. (B, C) SAG levels in 25-d-old plants WT and OX-*SNIPER1* plants treated with H_2_O (mock), 1 μM nlp20 (B) or 1 μM flg22 (C). Samples were collected for SAG measurement 24 hr after 1μM nlp20, or 9 hr after 1μM flg22 treatment. Different letters indicate genotypes with statistical differences (*p* < 0.05, one-way ANOVA test; n=3). All the experiments were repeated three times with similar results.

**Figure S2.**
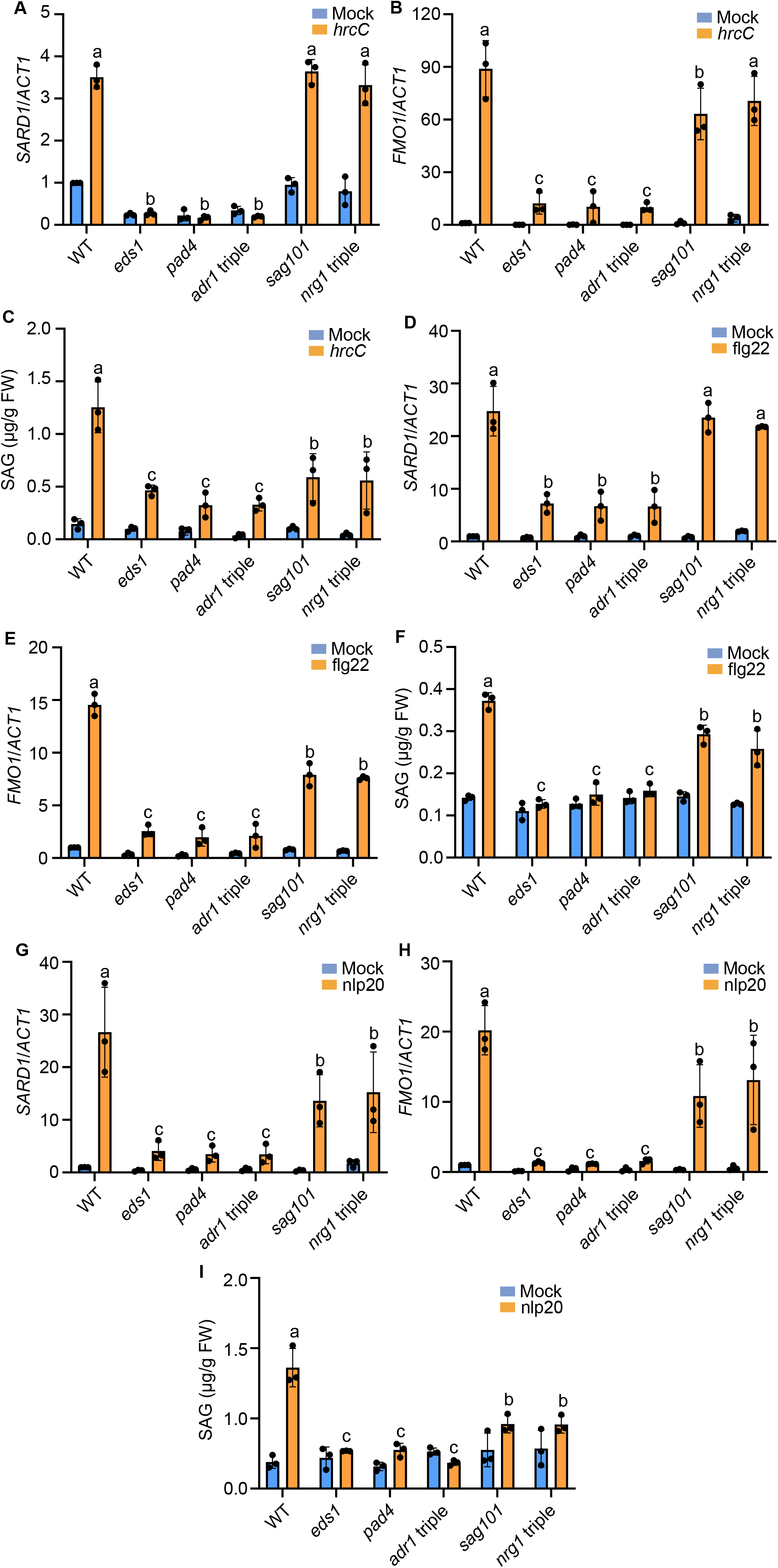
Induction of *SARD1* and *FMO1* gene expression and SAG production in TIR signaling mutants upon *Pto* DC3000 *hrcC*, flg22 or nlp20 treatment. (A, B) Relative expression levels of *SARD1* (A) and *FMO1* (B) in WT, *eds1-24, pad4-1, sag101-1, adr1 adr-L1 adr-L2* (*adr1 triple*) and *nrg1a nrg1b nrg1c* (*nrg1 triple*) mutant plants after treatment with *Pto* DC3000 *hrcC* (OD_600_ = 0.05) for 12 hours. Error bars represent SD from three different biological replicates. Different letters indicate genotypes with statistical differences (*p* < 0.05, one-way ANOVA test; n=3). (C, F, I) Four-week-old soil-grown plants of the indicated genotypes were treated with *Pto* DC3000 *hrcC* (OD_600_=0.05) (D), 1 μM flg22 (F) or 1 μM nlp20 (I). Samples were collected for SAG measurement 12 hr after inoculation of *Pto* DC3000 *hrcC*, 24 hr after treatment with 1 μM nlp20, or 9 hr after treatment with 1 μM flg22. Different letters indicate genotypes with statistical differences (*p* < 0.05, one-way ANOVA test; n=3). The experiment was repeated three times with similar results. (D, E) Relative expression levels of *SARD1* (D) and *FMO1* (E) in the indicated genotypes upon flg22 treatment. Total RNA was isolated from 12-d-old plate-grown seedlings 4 h after spraying with 1 μM flg22. Different letters indicate genotypes with statistical differences (*p* < 0.05, one-way ANOVA test; n=3). (G, H) Relative expression levels of *SARD1* (G) and *FMO1* (H) in the indicated genotypes upon nlp20 treatment. Total RNA was isolated from 12-d-old plate-grown seedlings 4 h after spraying with 1 μM nlp20. Different letters indicate genotypes with statistical differences (*p* < 0.05, one-way ANOVA test; n=3). For gene expression analysis in (A, B, D, E, G; H), the expression of each gene in the mock-treated WT plants was set as 1. All gene expression analyses were repeated twice with similar results.

**Figure S3.**
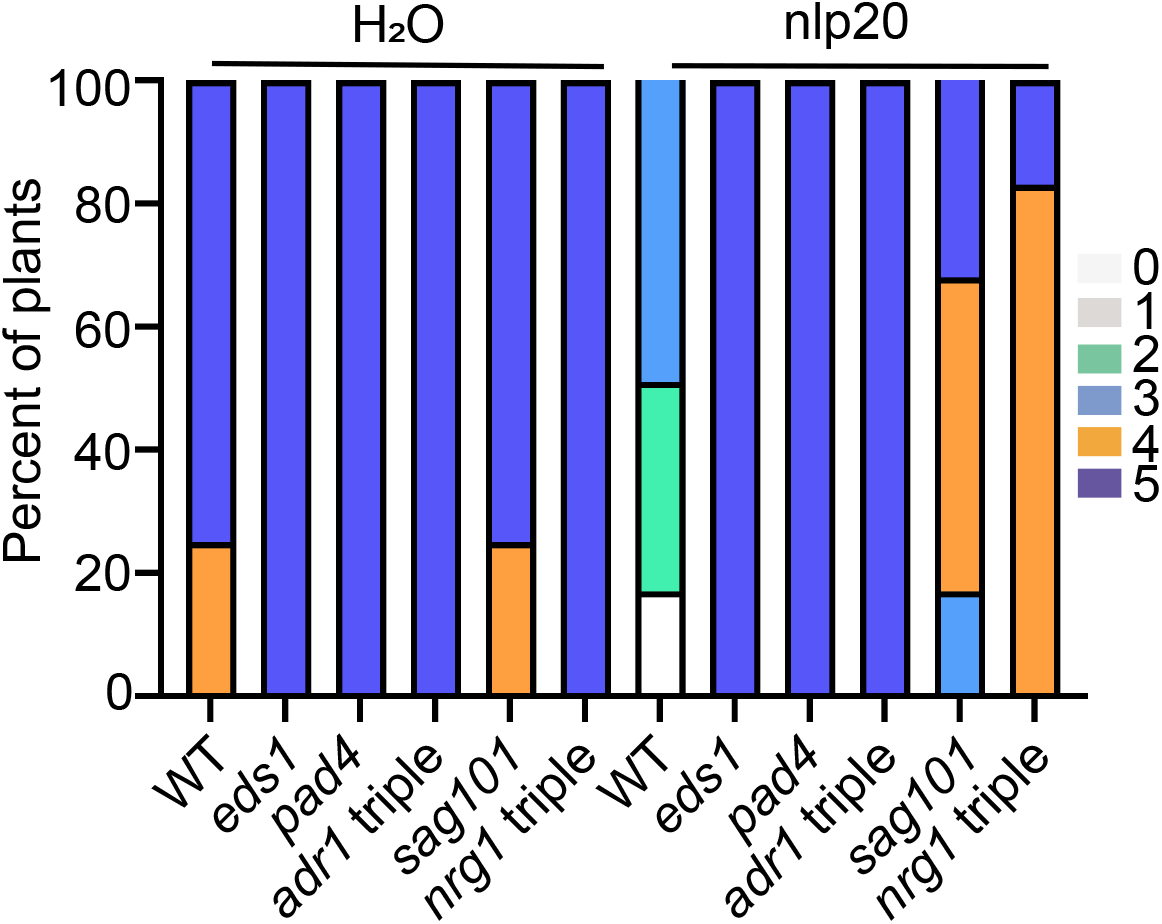
Growth of *Hpa* Noco2 on the distal leaves of TIR signaling mutants. mutants. Three-week-old soil-grown plants were pretreated with water or 1 μM nlp20 and sprayed with *Hpa* Noco2 spores (50,000 spores/ml) 24 hours later. Disease ratings are as described in Fig 1. The experiment was repeated twice with similar results.

**Figure S4.**
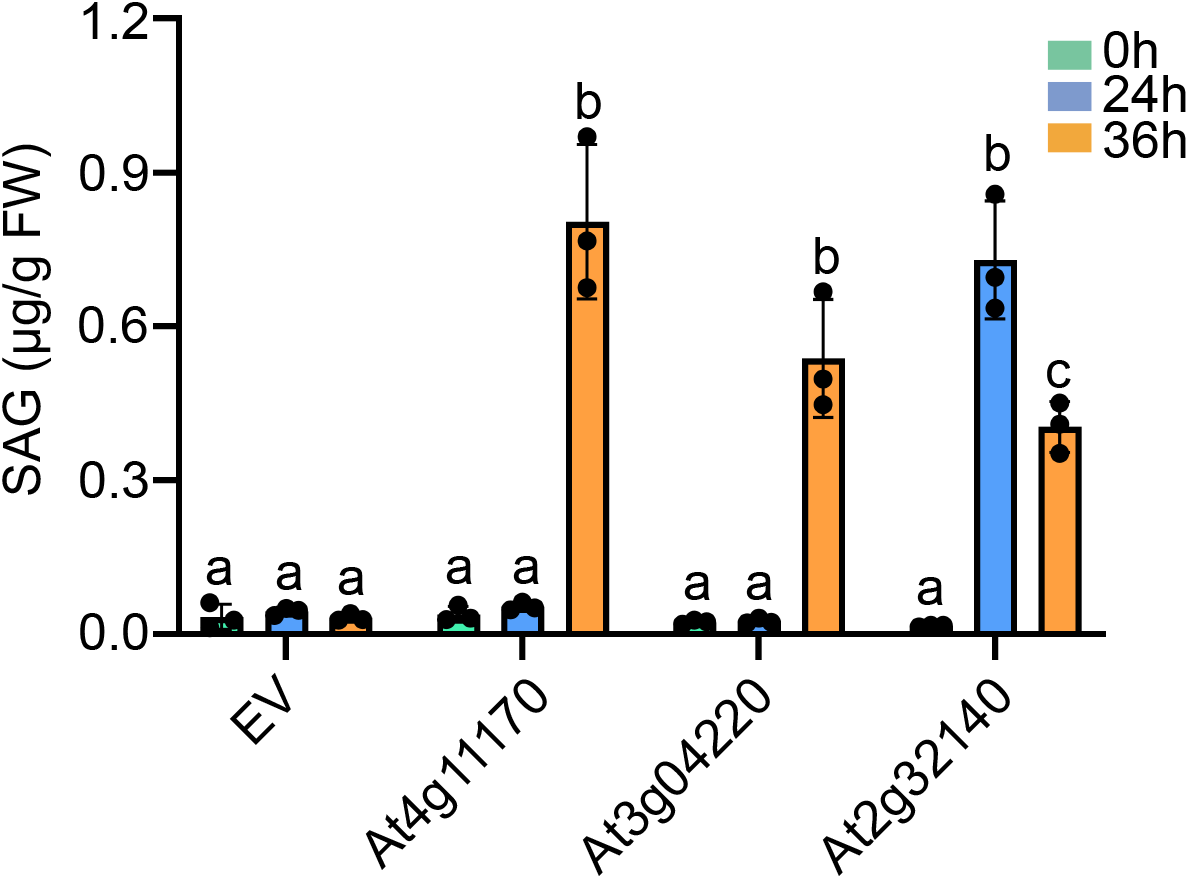
Levels of glucose-conjugated SA (SAG) in *N. benthamiana* leaves after infiltration of *Agrobacterium* carrying the *TIR* genes. Samples were collected 24 and 36 hours post infiltration of the bacteria (OD_600_=0.4) before HR was visible. Different letters indicate statistical differences (*p* < 0.05, one-way ANOVA test; n=3) compared with the empty vector control. The experiment was repeated twice with similar results.

**Figure S5.**
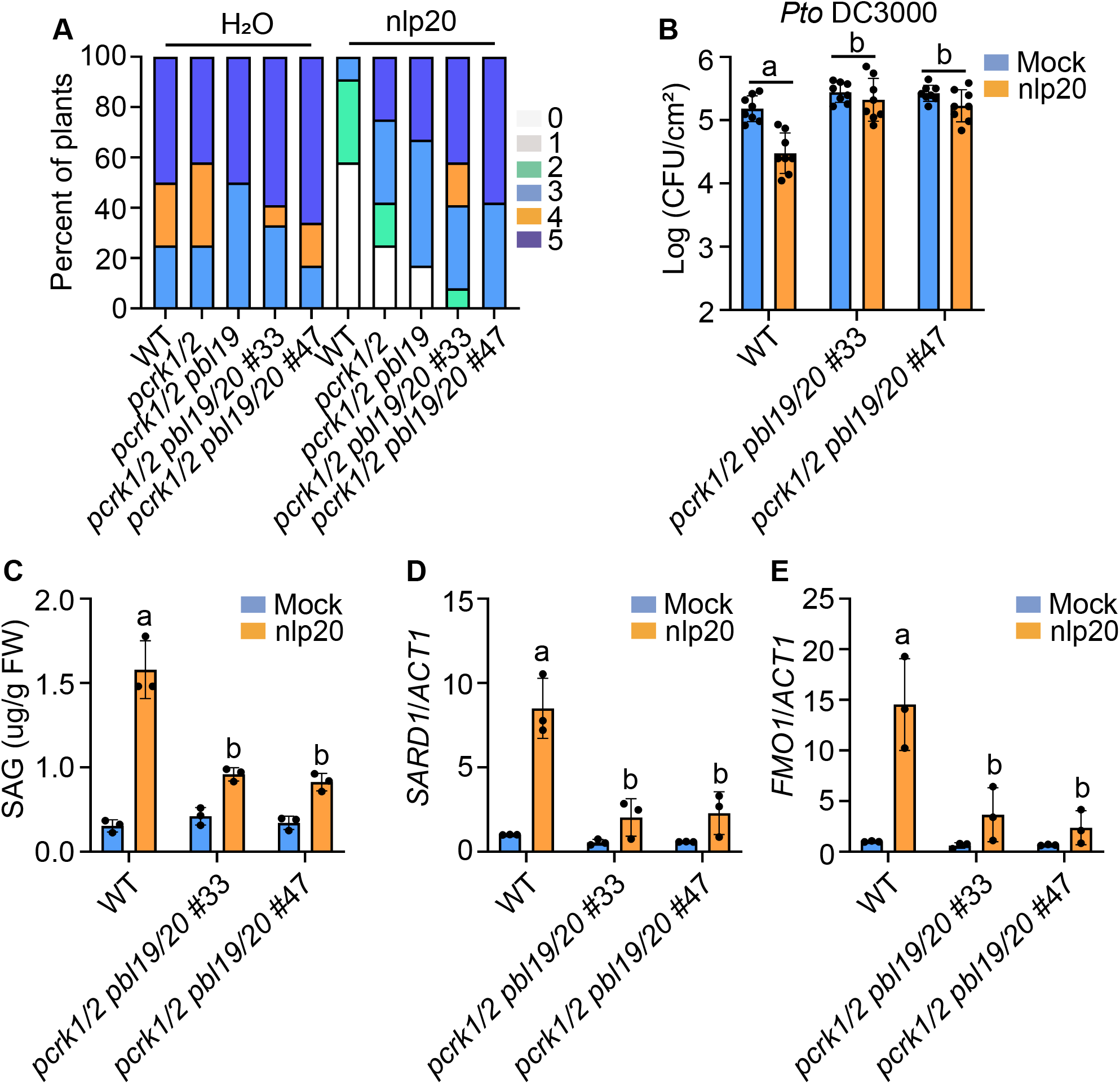
nlp20-induced immune responses are compromised in *pcrk1/2 pbl19/20* quadruple mutant plants. (A) Growth of *Hpa* Noco2 on the local leaves of WT, *pcrk1/2, pcrk1/2 pbl19, pcrk1/2 pbl19/20* #33 and *pcrk1/2 pbl19/20* #47 quadruple mutant plants after 1 μM nlp20 treatment. Infection was scored 7 dpi by counting the number of conidiophores per infected leaf. Detailed methodology was described in Fig 2E. (B) Growth of *Pto* DC3000 in the leaves of four-week-old WT, *pcrk1/2pbl19/20* #33 and *pcrk1/2 pbl19/20* #47 quadruple mutant plants pre-treated with water or 1 μM nlp20. 24 hours post elicitor treatment, the treated leaves were infiltrated with *Pto* DC3000 (OD_600_=0.001). Samples were taken 3 days after *Pto* DC3000 inoculation. Error bars represent SD from six biological replicates. The nlp20-induced protection among different genotypes was compared using two-way ANOVA test, and different letters indicate genotypes with statistical differences (*p* < 0.05, *n*= 6). (C) Levels of SAG in four-week-old soil-grown plants of the indicated genotypes treated with water or 1 μM nlp20. Samples were collected 24 hours post elicitor treatment. Different letters indicate statistical differences (*p* < 0.05, one-way ANOVA test; n=3) (D, E) Relative expression levels of *SARD1* (D) and *FMO1* (E) in the indicated genotypes. Total RNA extracted from 12-d-old plate-grown plants treated with 1 μM nlp20 for 4h. *ACT1* was used for normalization, and the expression of each gene in mock-treated WT was set as 1. Error bars represent SD from three different biological replicates. Different letters indicate genotypes with statistical differences (*p* < 0.05, one-way ANOVA test; n=3). All the experiments were repeated twice with similar results.

**Figure S6.**
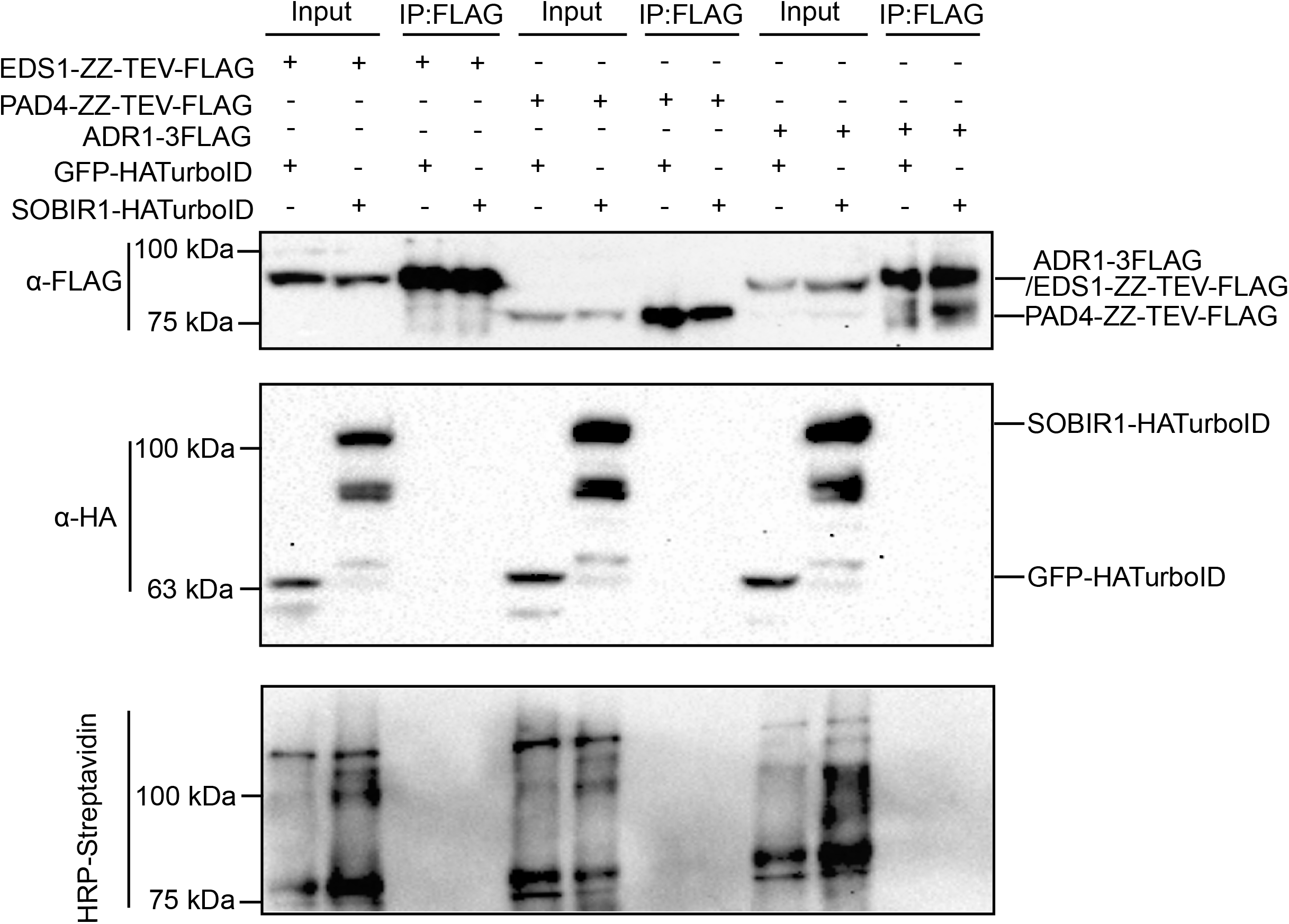
Analysis of interactions between SOBIR1 and EDS1/PAD4/ADR1 by TurboID and co-immunoprecipitation analysis. *Agrobacterium* carrying the indicated constructs were infiltrated into *N. benthamiana* leaves for protein expression. Immunoprecipitation of EDS1-FLAG-ZZ, PAD4-FLAG-ZZ or ADR1-3FLAG was carried out with anti-FLAG beads. The 3FLAG-tagged and HATurboID fusion proteins were detected by western blot using an anti-FLAG or anti-HA antibody by western blot. The biotinylated proteins were detected by western blot using HRP-Streptavidin. Molecular mass marker in kiloDaltons is indicated on the left. The experiment was repeated twice with similar results.

**Table S1.**
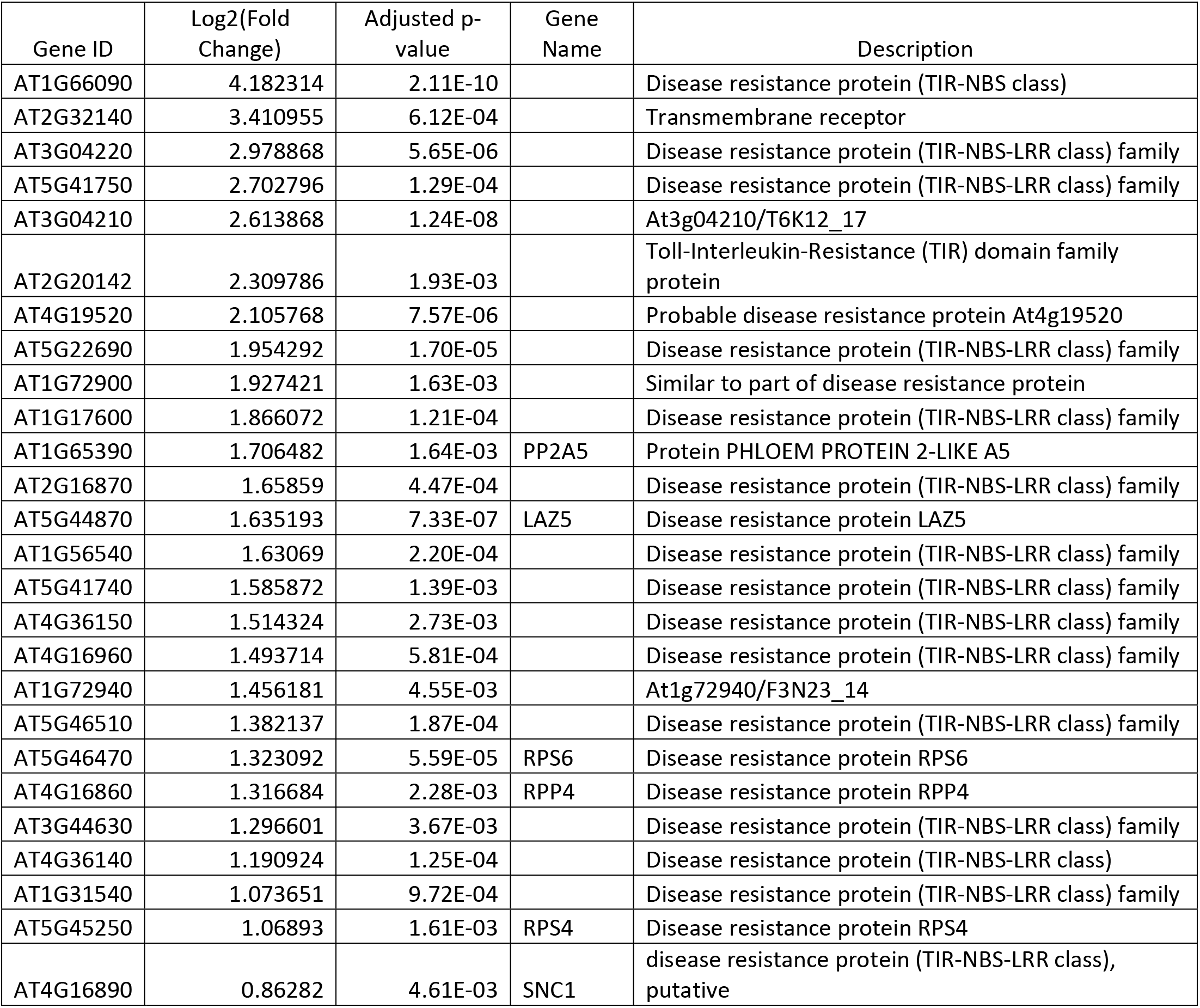
TIR-domain containing genes induced by nlp20 treatment for 1 hour compared with H_2_O treatment for 1 hour.

| Gene ID | Log2(Fold Change) | Adjusted p-value | Gene Name | Description |
| --- | --- | --- | --- | --- |
| AT1G66090 | 4.182314 | 2.11E-10 |  | Disease resistance protein (TIR-NBS class) |
| AT2G32140 | 3.410955 | 6.12E-04 |  | Transmembrane receptor |
| AT3G04220 | 2.978868 | 5.65E-06 |  | Disease resistance protein (TIR-NBS-LRR class) family |
| AT5G41750 | 2.702796 | 1.29E-04 |  | Disease resistance protein (TIR-NBS-LRR class) family |
| AT3G04210 | 2.613868 | 1.24E-08 |  | At3g04210/T6K12_17 |
| AT2G20142 | 2.309786 | 1.93E-03 |  | Toll-Interleukin-Resistance (TIR) domain family protein |
| AT4G19520 | 2.105768 | 7.57E-06 |  | Probable disease resistance protein At4g19520 |
| AT5G22690 | 1.954292 | 1.70E-05 |  | Disease resistance protein (TIR-NBS-LRR class) family |
| AT1G72900 | 1.927421 | 1.63E-03 |  | Similar to part of disease resistance protein |
| AT1G17600 | 1.866072 | 1.21E-04 |  | Disease resistance protein (TIR-NBS-LRR class) family |
| AT1G65390 | 1.706482 | 1.64E-03 | PP2A5 | Protein PHLOEM PROTEIN 2-LIKE A5 |
| AT2G16870 | 1.65859 | 4.47E-04 |  | Disease resistance protein (TIR-NBS-LRR class) family |
| AT5G44870 | 1.635193 | 7.33E-07 | LAZ5 | Disease resistance protein LAZ5 |
| AT1G56540 | 1.63069 | 2.20E-04 |  | Disease resistance protein (TIR-NBS-LRR class) family |
| AT5G41740 | 1.585872 | 1.39E-03 |  | Disease resistance protein (TIR-NBS-LRR class) family |
| AT4G36150 | 1.514324 | 2.73E-03 |  | Disease resistance protein (TIR-NBS-LRR class) family |
| AT4G16960 | 1.493714 | 5.81E-04 |  | Disease resistance protein (TIR-NBS-LRR class) family |
| AT1G72940 | 1.456181 | 4.55E-03 |  | At1g72940/F3N23_14 |
| AT5G46510 | 1.382137 | 1.87E-04 |  | Disease resistance protein (TIR-NBS-LRR class) family |
| AT5G46470 | 1.323092 | 5.59E-05 | RPS6 | Disease resistance protein RPS6 |
| AT4G16860 | 1.316684 | 2.28E-03 | RPP4 | Disease resistance protein RPP4 |
| AT3G44630 | 1.296601 | 3.67E-03 |  | Disease resistance protein (TIR-NBS-LRR class) family |
| AT4G36140 | 1.190924 | 1.25E-04 |  | Disease resistance protein (TIR-NBS-LRR class) |
| AT1G31540 | 1.073651 | 9.72E-04 |  | Disease resistance protein (TIR-NBS-LRR class) family |
| AT5G45250 | 1.06893 | 1.61E-03 | RPS4 | Disease resistance protein RPS4 |
| AT4G16890 | 0.86282 | 4.61E-03 | SNC1 | disease resistance protein (TIR-NBS-LRR class), putative |

**Table S2.** TIR-domain containing genes induced by nlp20 treatment for 6 hours compared with H_2_O treatment for 6 hours.

| Gene ID | Log2(Fold Change) | Adjusted p-value | Gene Name | Description |
| --- | --- | --- | --- | --- |
| AT1G57630 | 7.14555 | 3.84E-06 |  | Disease resistance protein RPP1-WsB, putative |
| AT4G11170 | 4.73465 | 4.44E-04 |  | Putative disease resistance protein At4g11170 |
| AT5G45090 | 3.91362 | 1.51E-02 | PP2A7 | Uncharacterized protein PHLOEM PROTEIN 2-LIKE A7 |
| AT1G47370 | 3.62641 | 2.83E-02 |  | Toll-Interleukin-Resistance (TIR) domain family protein |
| AT2G32140 | 3.59523 | 1.72E-02 |  | Transmembrane receptor |
| AT5G41750 | 3.33276 | 2.23E-04 |  | Disease resistance protein (TIR-NBS-LRR class) family |
| AT1G66090 | 3.02848 | 4.22E-04 |  | Disease resistance protein (TIR-NBS class) |
| AT1G72920 | 2.84247 | 3.10E-03 |  | Similar to part of disease resistance protein |
| AT5G45000 | 2.74362 | 1.49E-02 |  | Disease resistance protein (TIR-NBS-LRR class) family |
| AT2G20142 | 2.4793 | 2.51E-02 |  | Toll-Interleukin-Resistance (TIR) domain family protein |
| AT3G04220 | 2.28377 | 2.68E-02 |  | Disease resistance protein (TIR-NBS-LRR class) family |
| AT5G58120 | 1.90423 | 9.43E-03 |  | Disease resistance protein (TIR-NBS-LRR class) family |
| AT5G41740 | 1.71984 | 1.75E-02 |  | Disease resistance protein (TIR-NBS-LRR class) family |
| AT1G72900 | 1.71688 | 8.78E-02 |  | Similar to part of disease resistance protein |

**Table S3.**
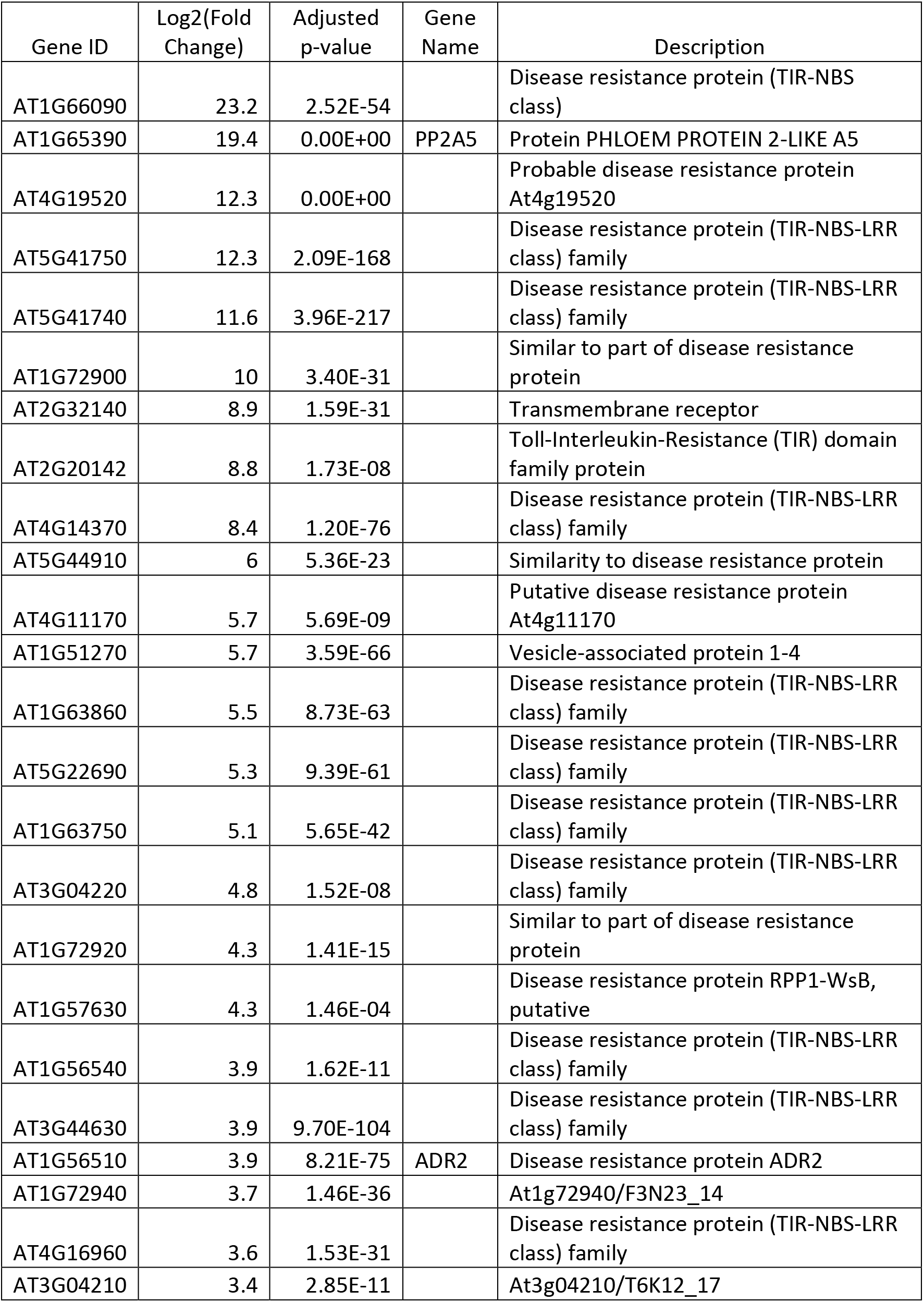
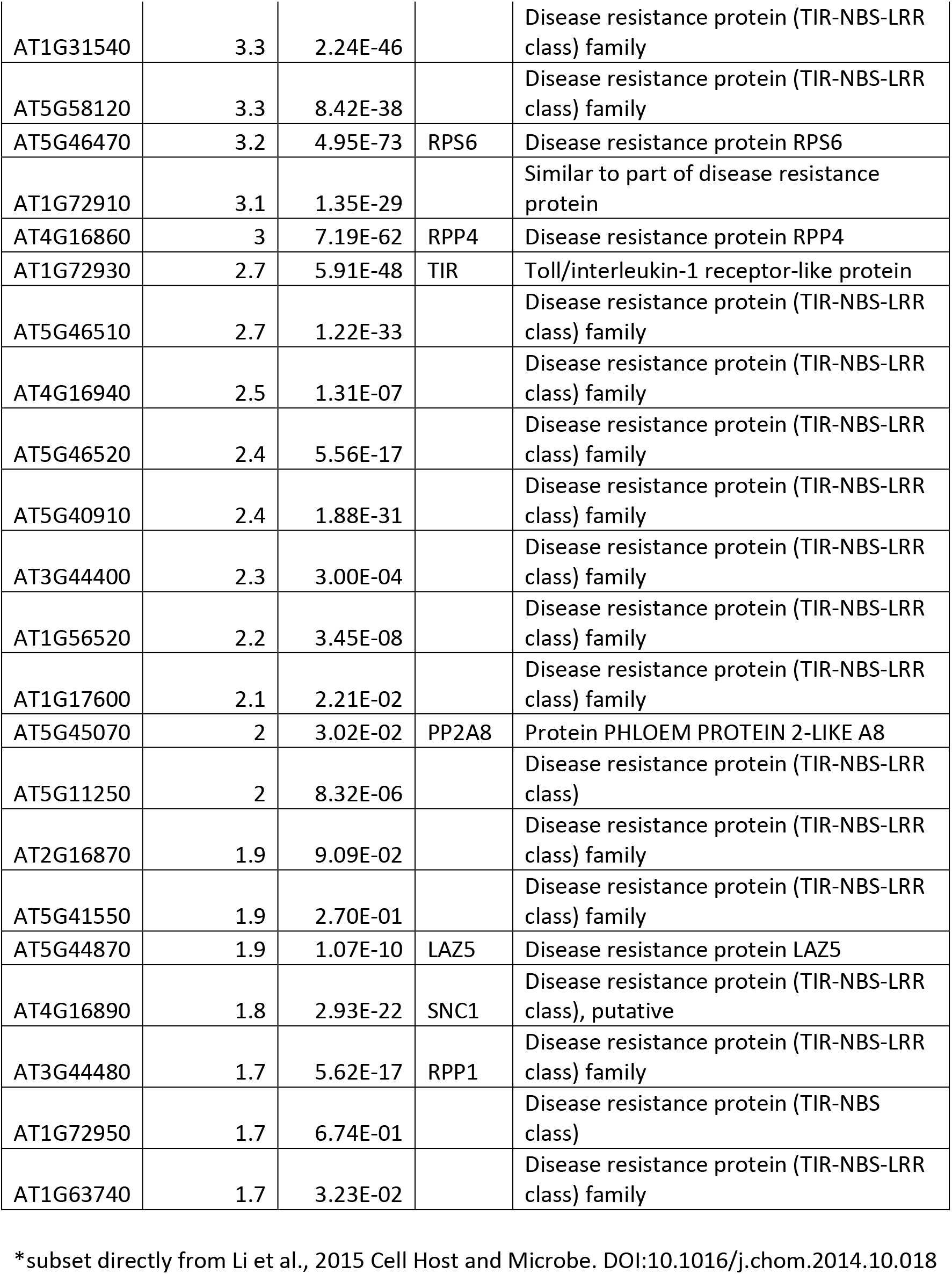
TIR-domain containing genes induced by flg22 treatment for 30 min compared with untreated*.

**Table S4.**
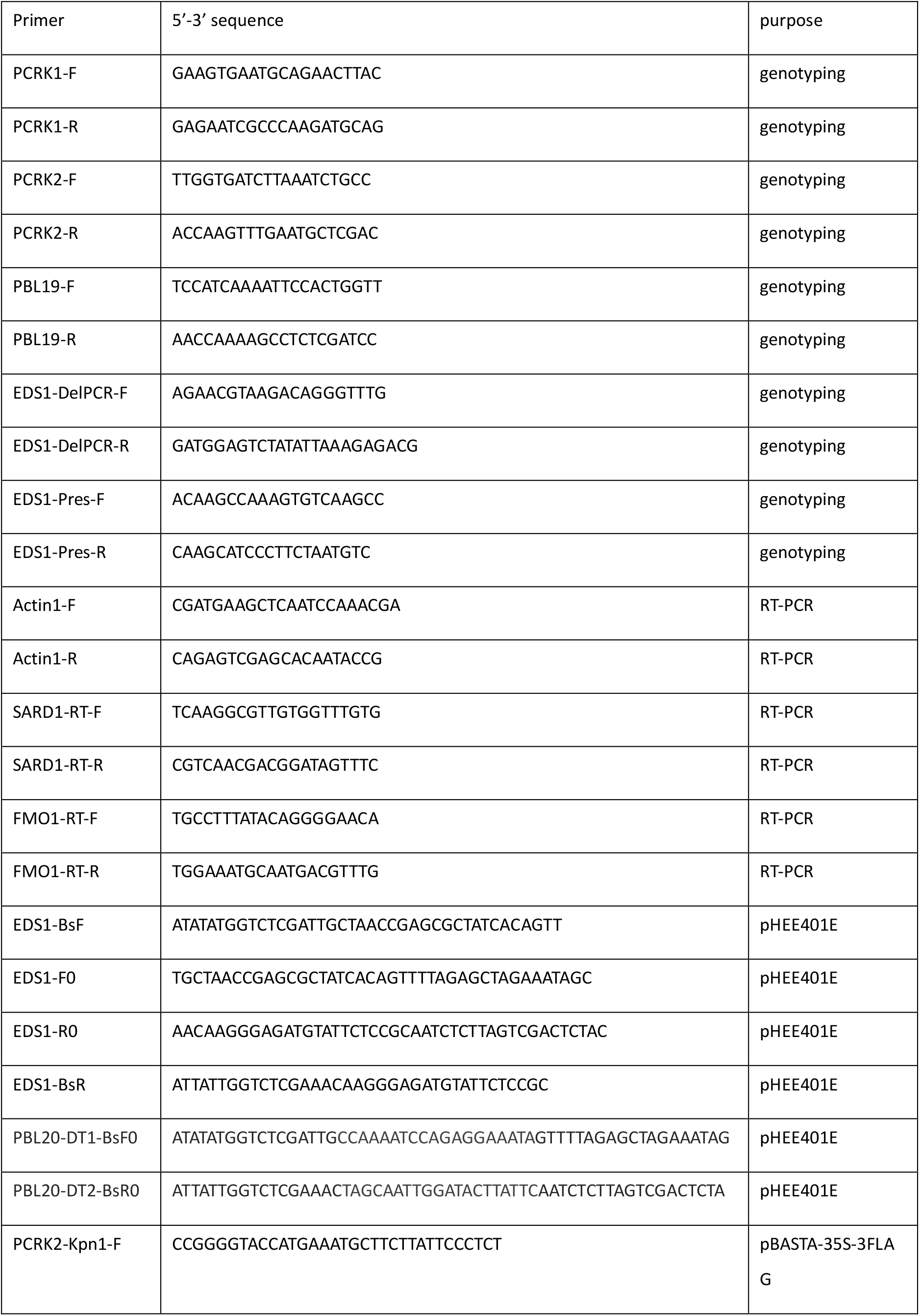
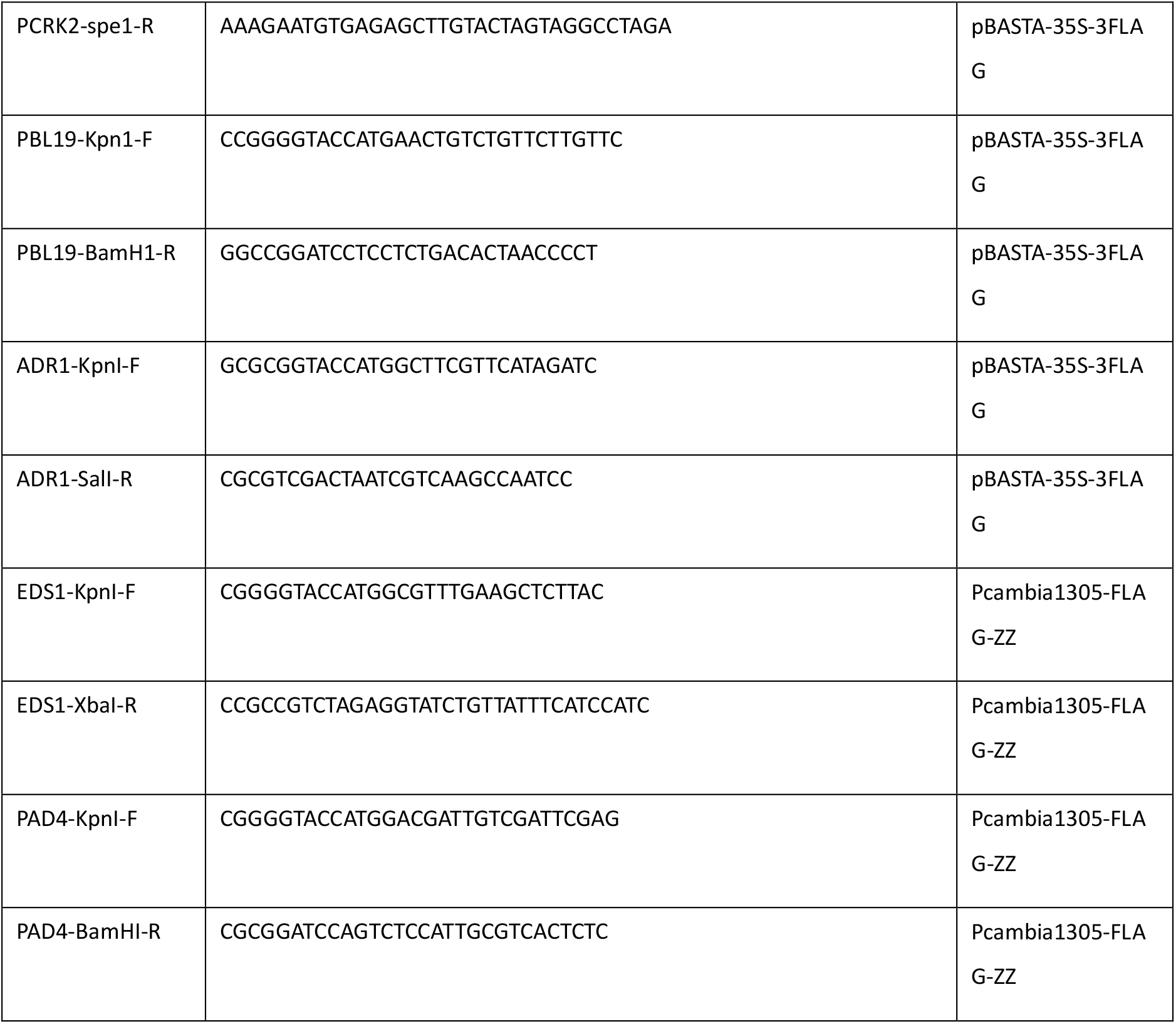
Sequences of primers used in this study.

## Notes

### Competing Interest Statement

The authors have declared no competing interest.

